# Foundation model for efficient biological discovery in single-molecule data

**DOI:** 10.1101/2024.08.26.609721

**Authors:** Jieming Li, Leyou Zhang, Alexander Johnson-Buck, Nils G. Walter

## Abstract

Modern data-intensive techniques offer ever deeper insights into biology, but render the process of discovery increasingly complex. For example, exploiting the unique ability of single-molecule fluorescence microscopy (SMFM)^1–5^. to uncover rare but critical intermediates often demands manual inspection of time traces and iterative *ad hoc* approaches that are difficult to systematize. To facilitate systematic and efficient discovery from SMFM data, we introduce META-SiM, a transformer-based foundation model pre-trained on diverse SMFM analysis tasks. META-SiM achieves high performance—rivaling best-in-class algorithms—on a broad range of analysis tasks including trace selection, classification, segmentation, idealization, and stepwise photobleaching analysis. Additionally, the model produces high-dimensional embedding vectors that encapsulate detailed information about each trace, which the web-based META-SiM Projector (https://www.simol-projector.org) casts into lower-dimensional space for efficient whole-dataset visualization, labeling, comparison, and sharing. Combining this Projector with the objective metric of Local Shannon Entropy enables rapid identification of condition-specific behaviors, even if rare or subtle. As a result, by applying META-SiM to an existing single-molecule Förster resonance energy transfer (smFRET) dataset^6^, we discover a previously unobserved intermediate state in pre-mRNA splicing. META-SiM thus removes bottlenecks, improves objectivity, and both systematizes and accelerates biological discovery in complex single-molecule data.

## Introduction

In the main strength of single-molecule observation also lies its greatest challenge: while it can reveal transient intermediates and other rare behaviors that would be masked in ensemble measurements, identifying such rare states in datasets of thousands of single-molecule traces is often time- and labor-intensive. While initial population-level characterization through FRET efficiency histograms^7,8^ or transition occupancy density plots (TODPs)^9^ is informative, such methods still require careful trace curation, do not capture the entirety of information contained in single-molecule traces, and can still obscure, e.g., minority FRET states or changes in kinetics through the underlying ensemble averaging. This is particularly true in multi-step biological processes such as translation and pre-mRNA splicing, where it may be impossible to strongly enrich for specific intermediates while maintaining conditions that closely resemble the native intracellular context. As a result, manual inspection of traces is still an indispensable step in many SMFM studies, introducing potential bias and slowing the pace of discovery. Furthermore, distinguishing biologically significant minority behaviors from artifacts is not trivial^10^, and often requires deep domain expertise in single-molecule biophysics. Thus, although the increased commercial availability of turnkey fluorescence microscopes has rendered SMFM more accessible, the *analysis* of single-molecule results continues to require extensive domain knowledge, and remains time- and labor-intensive despite the community’s progress towards standardizing and streamlining workflows^11–14^.

A key obstacle to devising a more comprehensive, systematic method for analysis and discovery in SMFM experiments is the diverse array of information it must encompass, such as (i) FRET efficiency as a measure of distances^7,8^; (ii) kinetics of state transitions^15–20^; (iii) the presence and number of photobleaching steps^21–23^; and (iv) the presence and kinetics of fluorophore blinking^24,25^. Extraction of this information is typically achieved through explicit physical or statistical models with well-defined parameters^26^ such as with Hidden Markov modeling (HMM)^27–29^. However, HMM idealizations reflect only a subset of the information contained in the data, potentially discarding useful information (e.g., non-Markovian behavior, or rapid dynamics that may superficially appear as measurement noise in an SMFM trace^15^). Furthermore, critical decisions such as which segments of traces to include in further analysis depend on features that are not straightforward to encapsulate in conventional algorithms, such as the sequence of disappearance of donor or acceptor fluorophores, the presence or absence of one or more photobleaching steps, and absence of intrinsic measurement artifacts. Due to these complexities and their varying applicability to different systems, researchers often develop customized tools only applicable to specific data analysis tasks and apply them in an *ad hoc* fashion, reducing the generality as well as the accessibility of these tools to non-experts^3^.

To address these challenges, we introduce a foundation model, META-SiM or **M**ultitask **E**nabled **T**ransferrable **A**ttention model neural network for processing **Si**ngle-**M**olecule fluorescence traces (Fig. 1). Rather than the task-specific training of earlier deep learning approaches^30,31^, META-SiM adopts a holistic approach, learning diverse characteristics of fluorescence traces in a network training strategy increasingly popular in natural language processing^32–34^, but underexploited in scientific research. This strategy divides the network training into upstream pre-training and downstream use cases with specific fine-tuning. Here, we pre-trained META-SiM with simulated fluorescence traces designed to capture a broad range of characteristics (Fig. 1b), and fine-tuned it with experimental or simulated fluorescence traces for specific downstream use cases (Fig. 1h, i, j), covering most single-molecule data analysis needs for one- and two-color measurements, including: (i) trace classification and segmentation; (ii) trace idealization; (iii) photobleaching step counting; and (iv) kinetic fingerprinting. The use of simulated traces for pre-training is important because, unlike experimental data, it provides training datasets of defined ground truth and much larger (∼1 million) than those typically collected in single-molecule experiments (hundreds to thousands). META-SiM achieves high performance in all four prototypical use cases, either with no further training or with fine-tuning on very small datasets (comprising ∼100 traces), demonstrating its high adaptability compared to previous deep learning approaches^14,30,31^.

**Fig. 1.**
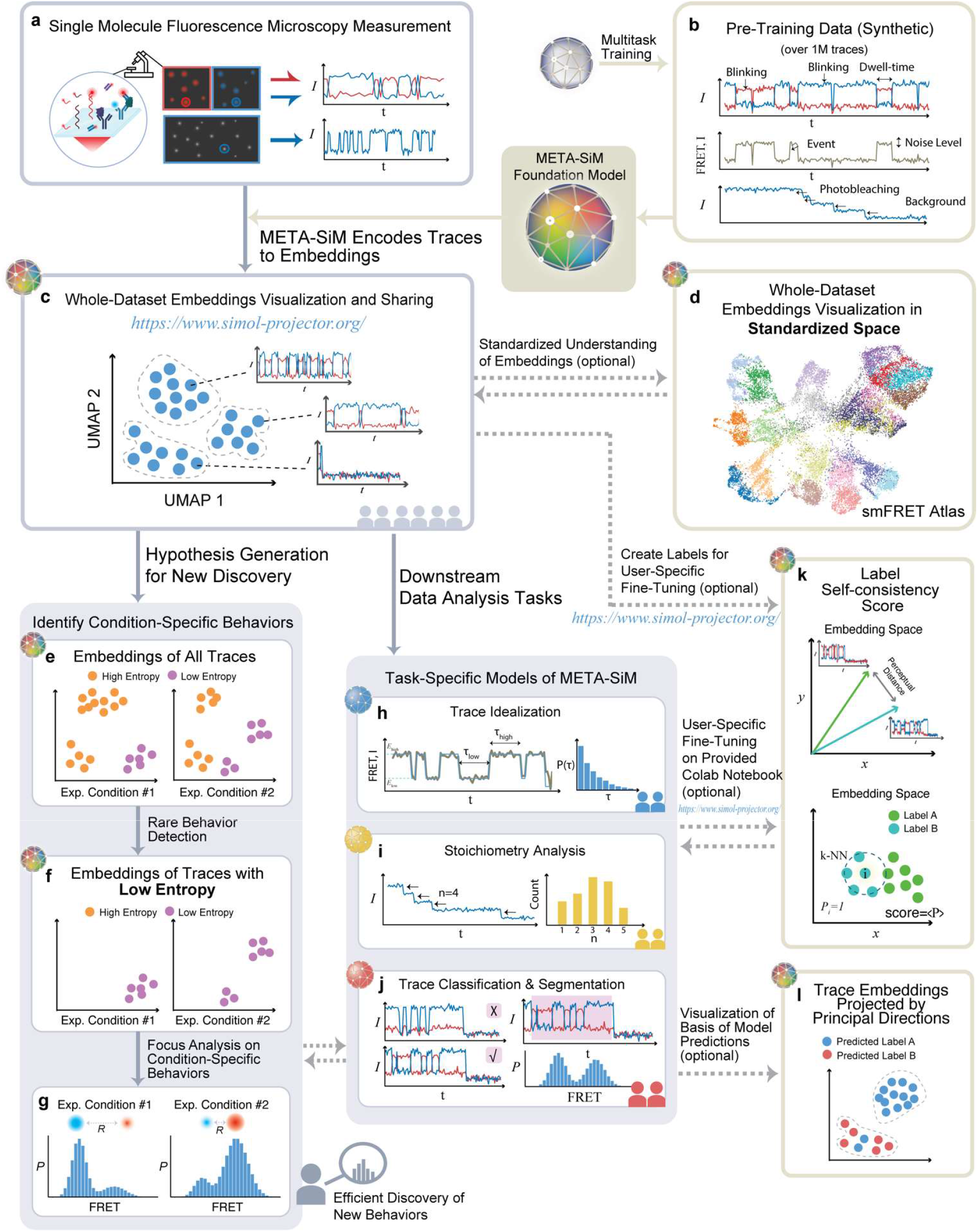
META-SiM enables analysis, visualization, and efficient discovery from diverse single-molecule datasets. **a**, Information from experimental time traces is encoded into high-dimensional embeddings by META-SiM for diverse downstream tasks. **b**, META-SiM is pre-trained with ∼1M synthetic traces on diverse common analysis tasks. **c**, The META-SiM Projector enables visualization of entire datasets based on information from the extracted embeddings. **d**, The embeddings from smFRET experiments can be projected into a global coordinate space, the smFRET Atlas, to facilitate comprehension and comparison. **e, f, g**, Local Shannon Entropy scores enable identification and analysis of rare or condition-specific behaviors, facilitating efficient discovery from complex datasets. **h, i, j**, META-SiM (with or without fine-tuning) can be used for diverse downstream analysis tasks. **k**, Optional manual labeling of a small subset (∼100 traces) of data from new types of experiments can be performed in the Projector, and assessed through a label Self-Consistency Score (SCS); these labels can be used to fine-tune the model for new tasks. **l**, Basis for model predictions visualized through Principal Projection.

Moreover, to aid in dataset visualization and exploration, we leveraged META-SiM’s unique ability to transform fluorescence time traces (Fig. 1a) into high-dimensional latent space embeddings to create the META-SiM Projector (Fig. 1c), an online interactive visualization tool that streamlines identifying patterns among complex single-molecule datasets. Using this Projector, we constructed an smFRET Atlas (Fig. 1d) as a step towards the comprehensive, standardized representation of information from smFRET traces in a common feature space. To conquer the “black-box” nature of deep neural networks, we introduce a Principal Projection function to help users interpret the clustering of projected traces and easily understand the model’s predictions (Fig. 1l). We also propose two useful metrics calculated from the META-SiM embeddings: a label Self-Consistency Score (SCS; Fig. 1k), which can be used to evaluate the quality of labels created by different users, as well as an objective method for training new users; and Local Shannon Entropy (LSE; Fig. 1e, f), which can be used as an index to quickly identify condition-specific behaviors, even when they are rare—often a key rate-limiting step in discovery. For instance, we show that projecting smFRET datasets from a previous study of paused transcriptional elongation complexes, color-coded by LSE, enables rapid identification of salt-dependent trace behaviors^35^.

Finally, as a more stringent test of the ability of META-SiM to expedite discovery in SMFM data, we applied it to an smFRET dataset from an *in vitro* study of the spliceosome in yeast whole cell extract^6^. Despite this system’s complex multi-state FRET dynamics, users with no prior knowledge of splicing exploited META-SiM and its LSE metric to efficiently (within one week) identify a new intermediate state, not previously detected by bioinformatic cluster analysis^6^, but later biochemically isolated^36^, describing an extended pre-mRNA conformation involved in 3’-splice site selection. These results illustrate the potential of META-SiM and its Projector to reduce the burdens of time and expertise, thereby accelerating the pace of discovery from complex single-molecule datasets.

### Multitask Pre-Training of META-SiM

As is common with AI foundation models,^37^ we employed a multitask training strategy (see Methods) in order to (1) reflect all essential capabilities in SMFM and (2) permit optimization of network hyperparameters to simultaneously accommodate all tasks. META-SiM comprises a multi-layer attention model that generates trace- and frame-level embedding vectors (Fig. 2), which are shared among tasks. For each task, we use a task-specific linear projector (“Head”) to convert the embeddings into the target space (Fig. 2). Pre-training was performed using 1 million synthetic traces that simulate a wide range of one- and two-color channel behaviors, including varying kinetics, numbers of states, and signal-to-noise ratio (SNR; Methods, Supplementary Table 1). While such synthetic data run the risk of exhibiting artificial traits or not representing the full range of possible experimental behaviors, leading to potential performance gaps when applied to experimental data, our multitask pretraining leaves no such gaps in downstream tasks (Fig. 3). We hypothesize that the diversity of our training tasks harmonizes the learning from individual tasks, allowing the model to learn fundamental traits more generalizable than those learned from only one or two specific tasks.

**Fig. 2.**
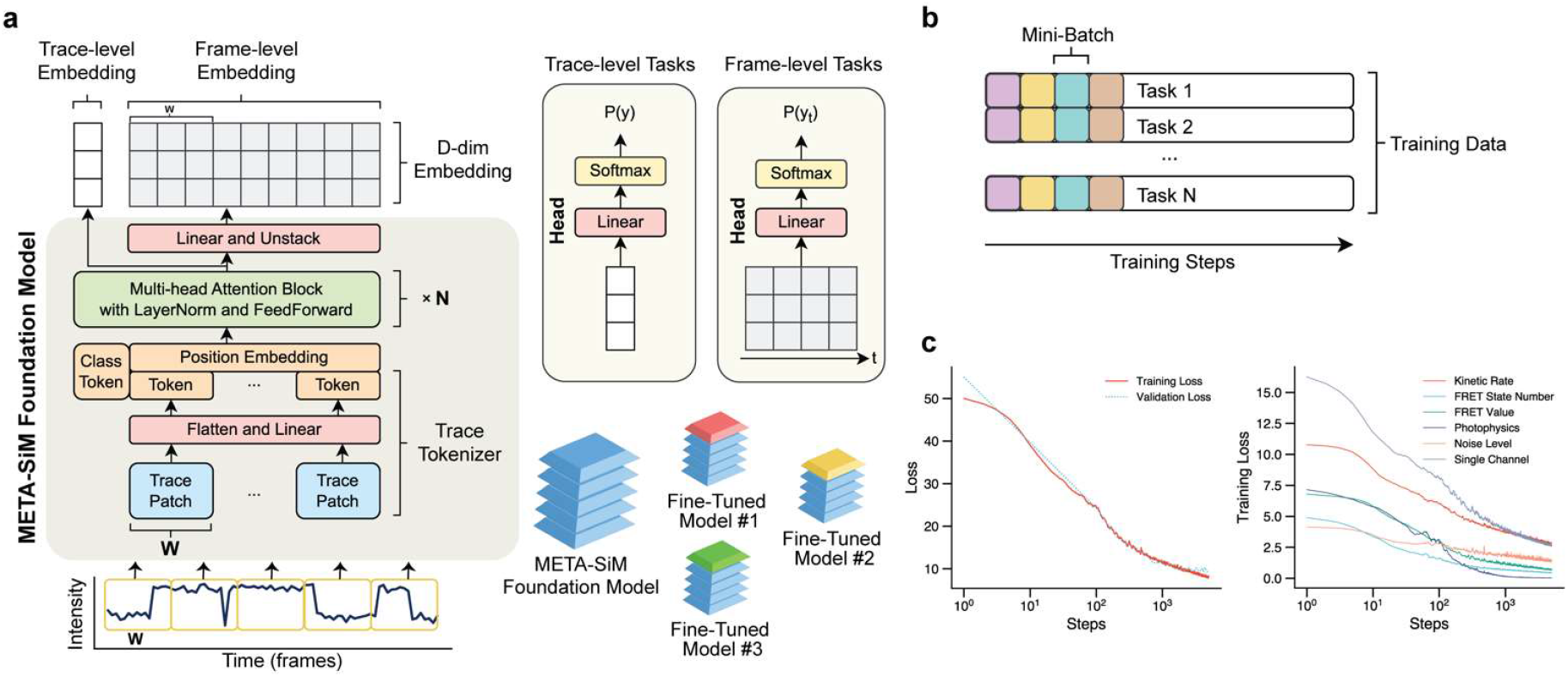
Architecture and training of META-SiM model. **a**, Architecture of META-SiM foundation model. An intensity *vs*. time trace of *n* frames is converted to *n*/*W* trace patches of width *W* frames. Trace patches are converted into tokens by a flatten operation, followed by a linear projection. The tokens are then combined with a position embedding, prepended by a class token, and fed into the transformer block with N multi-head attention layers, followed by a feed-forward network to create a trace-level embedding. A linear projection followed by an unstack operation transforms the output back to a D-dimensional frame-level embedding. For trace-level tasks, the trace-level embedding vector is used for linear logistic regression. For frame-level tasks, each frame-level embedding vector is used for linear logistic regression. **b**, Multitask training groups the simulated traces into mini batches with an equal split among training tasks to avoid catastrophic forgetting that can occur when tasks are trained sequentially. **c**, Loss function plots for the total loss (left) and for six representative tasks (right). All loss function values decreased, as expected, after 1,000 training steps, showing that the model learned from all the training task in parallel.

**Fig. 3.**
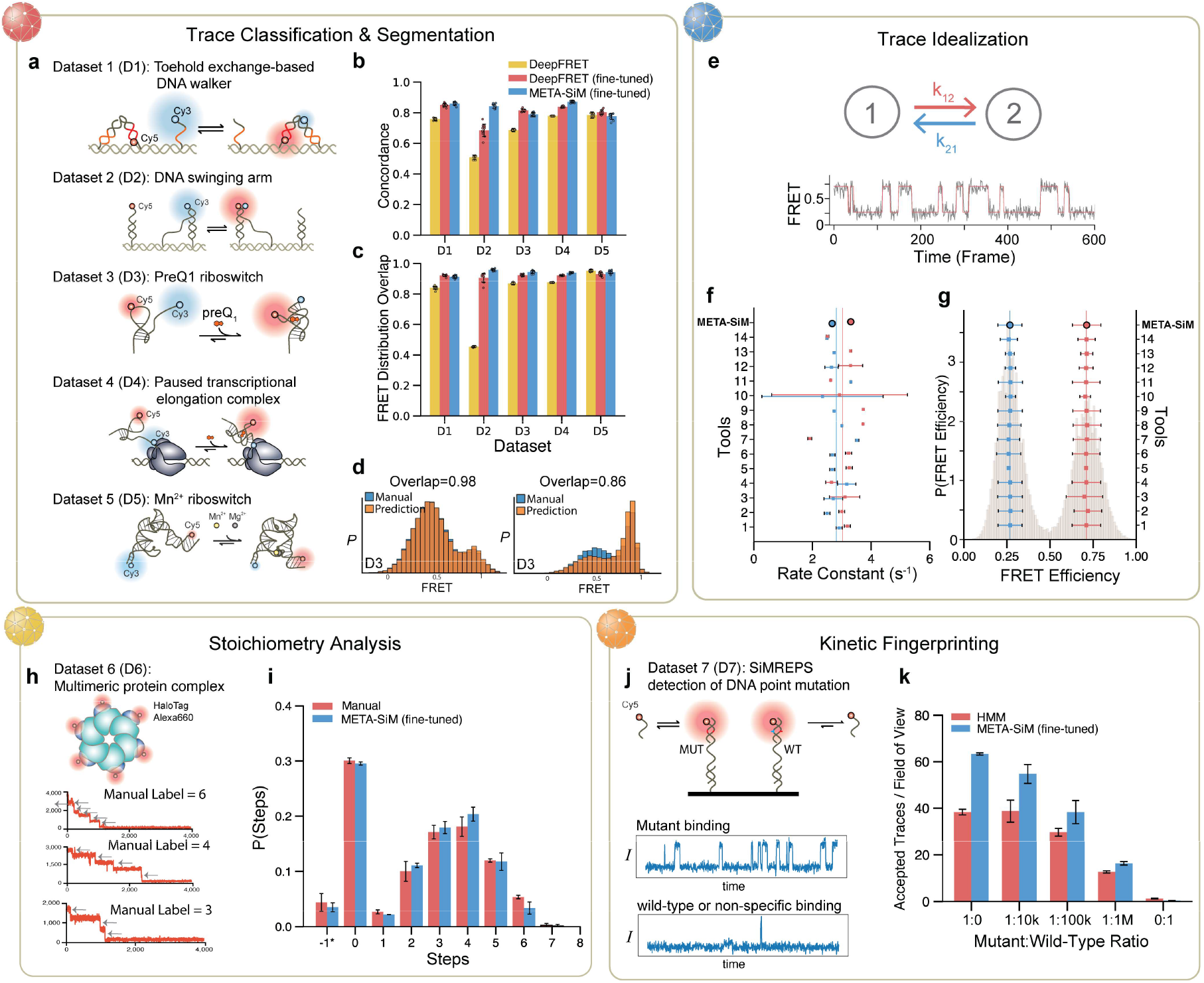
Performance of META-SiM on diverse analysis tasks. **a**,**b**,**c**,**d**, Evaluation of performance in trace segmentation and classification, and benchmarking against the recent AI model DeepFRET. **a**, Schematic of the five systems used in fine-tuning and testing: a toehold-exchange-based DNA walker (D1)^50^, a DNA swinging arm (D2)^51^, a preQ_1_ riboswitch (D3)^51^, a paused transcriptional elongation complex (D4)^52^, and a Mn^2+^ riboswitch (D5)^14^. **b**, The concordance of trace curation (acceptance or rejection for further analysis) of AI models with manual analysis. **c**, FRET distribution overlap between each AI model and manual analysis. **d**, Representative FRET histograms based on traces curated and segmented by META-SiM (fine-tuned) versus manual analysis (Supplementary Fig. 2b). **e**,**f**,**g**, Evaluation of performance in trace idealization for META-SiM not fine-tuned, and benchmarking against 14 other tools^15^: (1) Pomegranate; (2) Tracy(HMM); (3) FRETboard; (4) Hidden-Markury; (5) SMACKS(SS); (6) SMACKS; (7) Correlation; (8) Edge finding(CK); (9) Edge finding(k-means); (10) Step finding; (11) STaSI; (12) MASH-FRET(bootstrap); (13) MASH-FRET(prob); (14) postFRET. Time traces from a two-state FRET system (**e**) were idealized to estimate FRET efficiency in the two states (**g**) and create dwell time distributions (Supplementary Fig. 2c) that are fit with single exponential distributions to yield transition rate constants (**f**). **h, i**, Evaluation of performance in analysis of stoichiometry using a photobleaching dataset from a 6-subunit protein complex bearing up to one HaloTag Alexa Fluor 660 (Alexa660) label per monomer (**h**). (**i**) The distribution of photobleaching steps by manual counting performed by 3 trained experts *vs*. predictions by META-SiM fine-tuned for photobleaching steps counting. **j**,**k**, Evaluation of performance in kinetic fingerprinting and benchmarking against HMM-based analysis, in a use case involving detection of the *EGFR* point mutation T790M in DNA (**j**) with representative traces arising from of mutant (MUT) and wild-type (WT) or non-specific binding in the experiments. (**k**) MUT-like traces per field of view accepted by META-SiM *vs*. conventional HMM analysis and thresholding.

### META-SiM achieves high performance in diverse downstream tasks with minimal fine-tuning

Performance evaluation and benchmarking is important to establish not only META-SiM’s reliability as an analysis tool, but the accuracy and relevance of the information contained in its output embeddings that are employed for visualization and discovery. We considered four distinct use cases reflecting common needs in analysis of single-molecule time traces—trace classification and segmentation, trace idealization, stoichiometry analysis, and kinetic fingerprinting—and benchmarked performance against existing state-of-the-art or commonly used tools (Fig. 3, Methods, Supplementary Fig. 1). In contrast to earlier task-specific neural network-based tools, META-SiM rivals both advanced computational tools and manual analysis on all tasks. The model successfully curated and segmented traces from five distinct smFRET datasets to build accurate FRET histograms (Fig. 3a-d); idealized traces in a two-state system to extract accurate dwell time distributions, rate constants, and FRET states (Fig. 3e-g); counted photobleaching steps in close agreement with manual analysis (Fig. 3h, i); and discriminated between a point mutant and wild-type DNA sequence by SiMREPS kinetic fingerprinting^38^ with higher sensitivity and specificity than the original HMM-based analysis (Fig. 3j, k). While supervised fine-tuning of the model based on user-labeled examples using a logistic regression (see Methods) was used to improve the accuracy of the model at most tasks, since such fine-tuning only involved re-training the final layer of the network, it could be achieved with only ∼100 examples, 10-to 100-fold fewer than for previous task-specific tools based on deep neural networks^14,30,31^, and could thus be performed locally on a standard laptop computer (see Methods). In addition, this fine-tuning can typically be skipped entirely if a desired task, such as trace idealization, is already among the tasks the model was pre-trained on (Fig. 3f, g, Supplementary Table 1).

### META-SiM Projector: a flexible and powerful tool for visualizing whole single-molecule datasets

The complexity and heterogeneity of single-molecule datasets makes them challenging to visualize in a comprehensive manner with conventional methods, frequently necessitating labor-intensive inspection of individual traces to develop a thorough understanding of an experiment. Since META-SiM generates a trace-level embedding vector broadly encoding the attributes important for the multiple analysis tasks on which the model was trained, entire single-molecule datasets (regardless of complexity) can be visualized by projecting these high-dimensional embeddings into a cloud of points in 2D or 3D space using dimensionality reduction algorithms such as PCA^39^, UMAP^40^ and t-SNE^41^, where each point represents a single time trace (Fig. 4a). To test whether the pattern of point clusters in these projections meaningfully represents the clustering of relevant data attributes, we constructed and visualized synthetic time trace datasets―not used for training―with controlled variations in commonly considered data attributes such as SNR (Fig. 4b), state dwell times (Fig. 4c), FRET values (Supplementary Fig. 2a), number of FRET states (Supplementary Fig. 2b), and number of photobleaching events (Supplementary Fig. 2c) while keeping other attributes constant. The clustering of traces with similar traits in the 2D UMAP plots aligns well with variations in each attribute, demonstrating the effectiveness of the dimensionality reduction. Furthermore, when we performed the UMAP operation on experimental traces from datasets D1 (Fig. 4d) and D2-D5 (Supplementary Fig. 2d-g) that were manually either accepted or rejected by researchers for further analysis, the clustering of points was correlated with the labeled subgroups (Fig. 4d). Note that the META-SiM Projector also allows individual traces to be inspected and their nearest-neighbor traces (most similar in embedding vector) to be highlighted by clicking on points in the projection, enabling interactive exploration of the structure of heterogeneous single-molecule datasets.

**Fig. 4.**
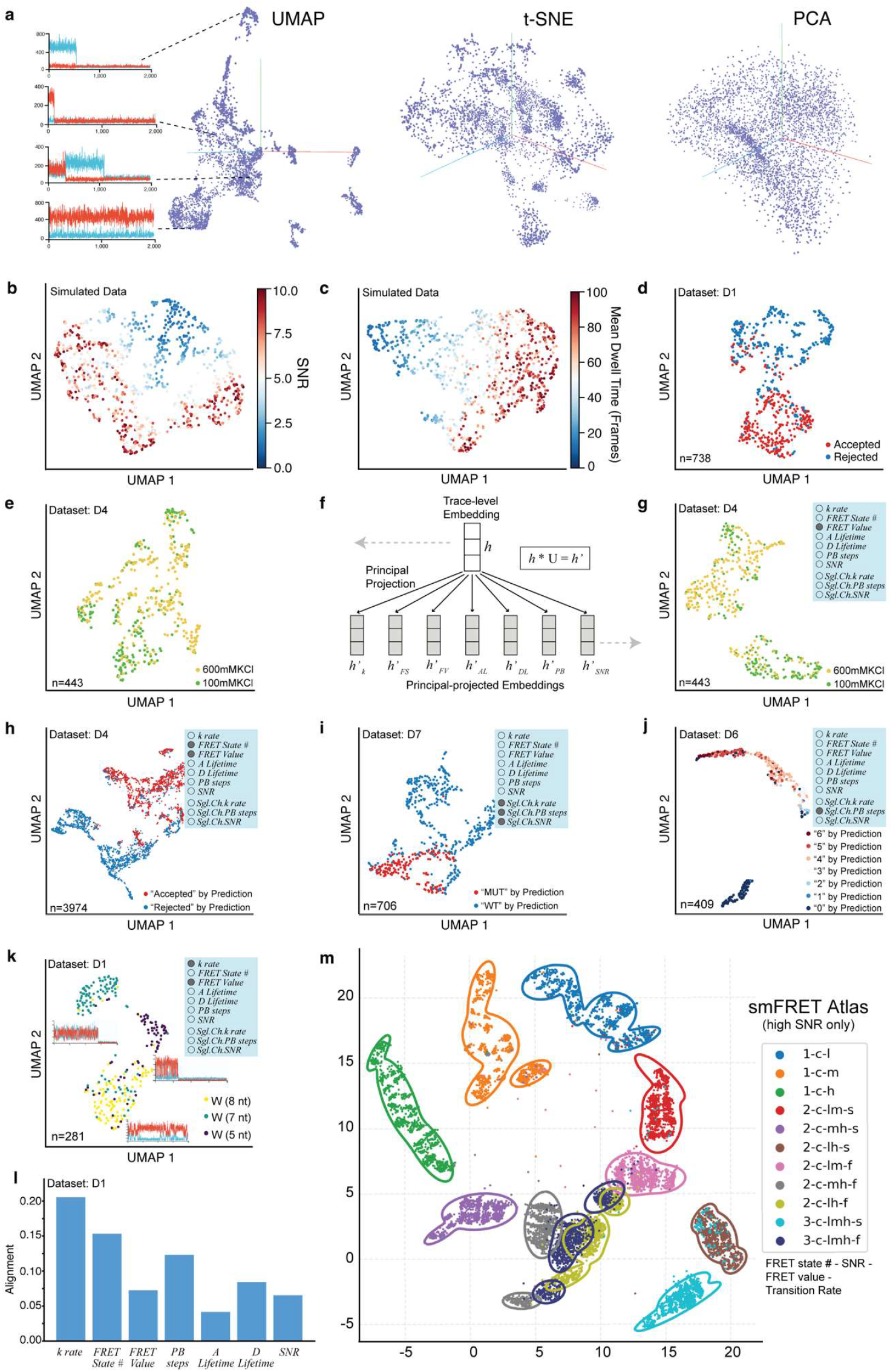
Whole-dataset visualization and interpretation with META-SiM Projector. **a**, 3D projections of high-dimensional embeddings for an smFRET dataset created with UMAP, PCA and t-SNE algorithms. Each point represents a projection of the high-dimensional embedding of a single time trace into 3D space. **b, c**, 2D UMAP projections of simulated traces with varying signal-to-noise ratio (SNR) (**b**) or mean dwell time (**c**). **d**, 2D UMAP projection of traces from dataset D1^35^ that were manually accepted (red) or rejected (blue) for further analysis. **e**, 2D UMAP projection of the traces from a dataset D4^6^ collected in 600 mM or 100 mM KCl. **f**, Schematic of the Principal Projection operation. The general dimensionality reduction uses the trace-level embedding *h* generated from META-SiM, while the Principal Projected embedding *h’* is generated by projecting *h* with a low-rank matrix U, which is generated by low-rank decomposition of the head layers grouped by task. **g**, 2D UMAP projection of the same traces in panel (**e**), but with Principal Projection by FRET Value. **h, i, j**, 2D UMAP projections of the traces from datasets D4 (**h**), D7 (**i**), and D6 (**j**) colored by the predicted labels from META-SiM classifiers, with Principal Projection according to different attributes of interest as indicated in the blue inset boxes. *k rate* = transition rate; *FRET State #* = number of FRET states; *FRET Value* = trace-wide FRET efficiency; *A Lifetime* = acceptor lifetime prior to bleaching; *D Lifetime* = donor lifetime prior to bleaching; *PB steps* = photobleaching steps; *SNR* = signal-to-noise ratio; *Sgl*.*Ch*.*k rate* = single-channel transition rate; *Sgl*.*Ch*.*PB steps* = single-channel photobleaching steps; *Sgl*.*Ch*.*SNR* = single-channel SNR. **k**, 2D UMAP projections of traces from dataset D1 where DNA walkers of different toehold lengths (8, 7, or 5 nucleotides, nt) yielded different stepping rates. The Principal Projection by kinetic rate and FRET value yields clustering of traces by toehold length, illustrating clear differences in behavior among these three constructs. **l**, Alignment of specific data analysis attributes with the META-SiM classifier. **m**, smFRET Atlas showing a UMAP projection of 22,000 simulated traces grouped by behavior, according to the following nomenclature: “FRET state number–SNR–FRET value–transition rate.” FRET state number takes values of “1”, “2”, or “3”; SNR takes values of “c” (clean, or high SNR) and “n” (noisy, or low SNR); FRET value takes values of “h” (high), “m” (middle), and/or “l” (low); transition rate takes values of “s” (slow) and “f” (fast). Contours indicate where the fitted probability density of the corresponding cluster drops to 0.01; a trace outside the contour is thus unlikely to belong to the corresponding cluster.

While these projections are comprehensive, a limitation is that they may not always yield strong clustering with respect to a variable of interest (Fig. 4e). To enable visualization of datasets based on specific data traits, we leveraged our multitask training strategy to group the 30 training task heads into seven attributes (three attributes for single-channel data; Supplementary Table 1), and calculated the operator for each attribute by low-rank matrix decomposition. In this way, an embedding projected to each attribute’s dimension was generated before the dimensionality reduction operation, yielding a “Principal Projection” that reflects the desired combination of traits (Fig. 4f, Methods). Thus, when specific data attributes are known or suspected to be important in differentiating populations of traces, this Principal Projection operation can be employed to more clearly segregate populations into meaningful clusters (Fig. 4g) than the general UMAP operation (Fig. 4e). In addition, the Principal Projection permits visualization of the basis of decisions by the network, such as those made in classifying traces or counting photobleaching steps (Fig. 4h-j). The network-predicted labels are clustered differently under different Principal Projections (Supplementary Fig. 3), providing insights into the most dominant differentiating factor(s) in the network’s decision-making process. The fractional contribution of each attribute in decision-making can be quantified as the correlation between the network’s head layer and each Principal Projection operator (see Methods), notably yielding quantitative insights into the importance of each attribute in predictions by the network (Fig. 4l, Supplementary Fig. 4). The Principal Projection can also yield tighter clustering based on known combinations of traits, providing a potential basis for multiplexed detection of distinct molecular species based on their trace characteristics. For example, three related macromolecular complexes (DNA walker systems with different stepping rates) were separated well in a Principal Projection based on two dimensions: kinetic rate and FRET value (Fig. 4k).

Although the above methods can greatly facilitate understanding of complex single-molecule datasets and classification decisions made by the network, the experiment-specific projections are not optimal for comparisons between experimental systems or laboratories. To establish a universal framework for such comparisons, we constructed a UMAP model based on 1 million simulated traces to create an smFRET Atlas capturing a broad range of commonly relevant characteristics (Fig. 4m): FRET state number, SNR, FRET values, and transition rates. In this Atlas, traces with similar characteristics (e.g., rapid 2-state transitions between a low- and a mid-FRET state with high SNR) cluster together, and the separation between these clusters increases when only high- or low-SNR traces are considered (Supplementary Fig. 5). This smFRET Atlas thus provides a common coordinate space for visualizing and comparing traces from different datasets based on their FRET states and dynamics, with the potential to standardize SMFM analysis.

### Model-derived metrics enable systematic quality control and hypothesis generation

The information META-SiM stores in embedding vectors provides a basis not only for analysis and visualization of single-molecule data, but for extracting quantitative metrics that assess the consistency of analysis or identify behaviors unique to certain experimental conditions. For example, consistency of trace labeling and segmentation—either for conventional analysis or fine-tuning of META-SiM—is a critical aspect of single-molecule analysis, but challenging to objectively assess due to the wide range of data characteristics and use cases that must be considered. To address this need, we leverage the detailed information about trace characteristics encapsulated in the high-dimensional embeddings to define label SCS, whose value (ranging from 0 to 1) indicates the degree of alignment with labels for similar traces (Figs. 1k, 5a; Methods). The label SCS empowers researchers to not only produce high-quality training data for the foundation model, but also to identify possible human errors during manual data analysis. For each of the five datasets examined here, the label SCS initially increases as a researcher labels an increasing fraction of traces, approaching an asymptotic consistency score (generally ≥ 0.8) after ∼5-20% of traces are labeled (Fig. 5b). The SCS varies among individual experiments for any given system, but the mean score is generally above 0.8 for our five experimental datasets (Fig. 5c). As many experiment-specific factors can impact labeling consistency, one should be cautious not to over-interpret the absolute value of the SCS between different experimental systems. However, for a given experimental system, the SCS provides meaningful quantitative feedback about labeling consistency. Based on these results, we suggest targeting a label SCS of at least 0.8 as a general rule-of-thumb (Fig. 5c); a lower score may signal inconsistent analysis and/or indicate that further labeling should be performed.

**Fig. 5.**
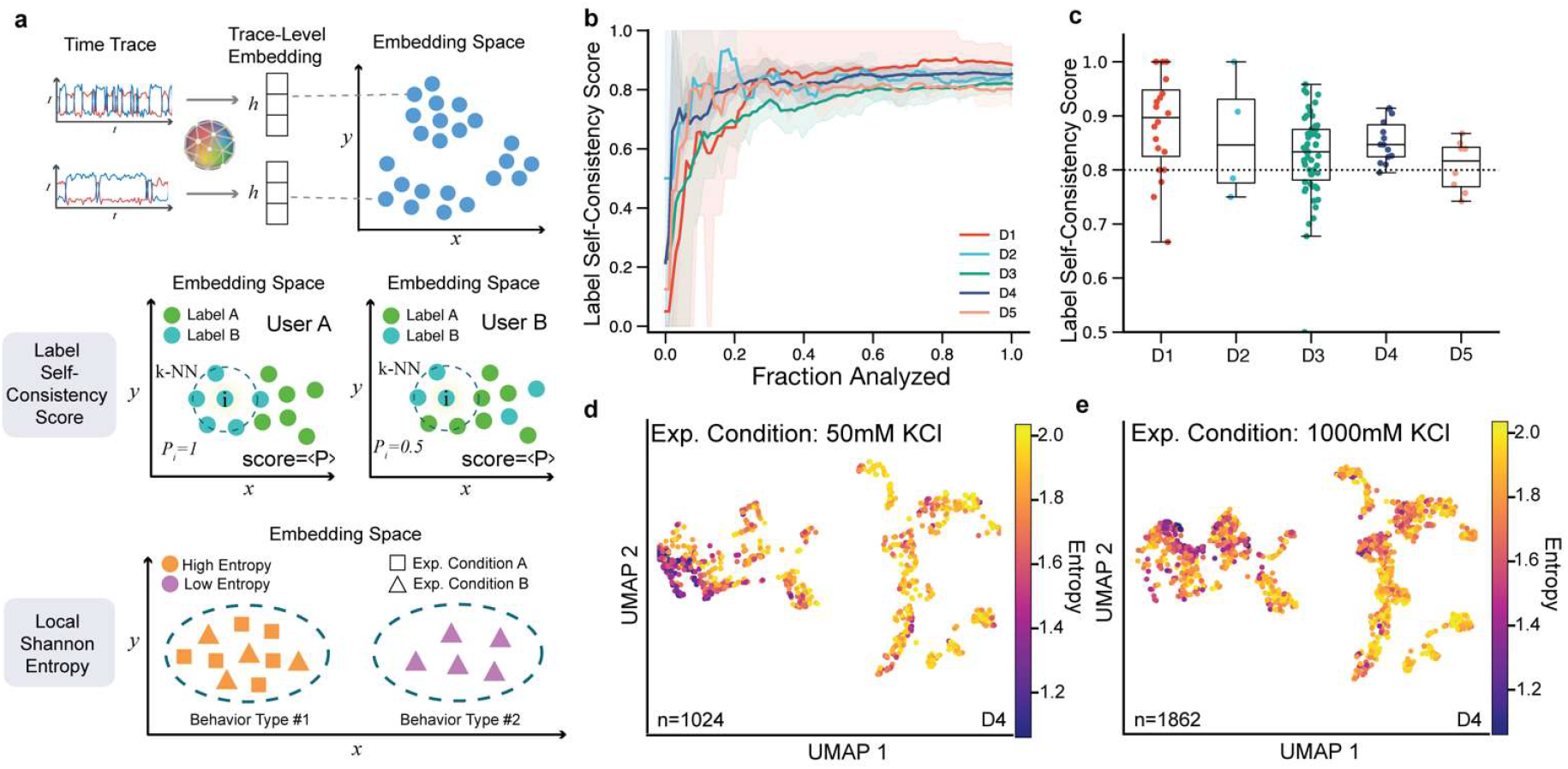
Quantitative metrics for quality control and discovery. **a**, Schematic of two proposed metrics: label Self-Consistency Score (SCS) and Local Shannon Entropy (LSE). The distance between two trace-level embeddings is used to quantify the similarity of any two traces, which is used as a basis for both metrics. The label SCS is a metric from 0 to 1 that is the average of the probability *P*_*i*_ in a dataset, where *P*_*i*_ is the probability of any given k-nearest neighbor (k-NN) trace, denoted by the dashed circle, having the same label as the center trace *i*. In this example, User A’s labeling has a higher score than User B’s since the labels within the area enclosed by the dashed line are more similar to that of trace *i*. LSE is a non-negative metric that quantifies the experiment condition complexity of the target trace and its k-NN traces by calculating the Shannon entropy of the probability distribution of the traces’ experiment conditions. A high entropy (Behavior Type #1) indicates that a behavior can be found in multiple datasets or experimental conditions (e.g., in Experimental Conditions A and B), and a low entropy (Behavior Type #2) indicates that a behavior can be found in fewer datasets or conditions (e.g., only in Experiment Condition B). **b**, Label SCS of individual users as a function of the fraction of traces analyzed. **c**, Label SCS distribution of movies in datasets D1-D5 from 5 individual users. **d, e**, UMAP projection of two experimental conditions from dataset D4 where each trace is colored by LSE. The low-entropy traces are clustered at different locations on the UMAP at different concentrations of KCl, indicating the presence of KCl-dependent behaviors in the corresponding smFRET traces.

Another critical challenge in single-molecule analysis is the identification of new or unique behaviors in large and heterogeneous datasets. This conventionally requires *ad hoc* approaches that involve an initial (often manual) trace curation, followed by population-level characterization by FRET histograms, TODPs, and/or kinetic analysis (e.g., dwell time or cross-correlation analysis) tailored to the system in question. Because it may not be known in advance what new behaviors will arise under certain experimental perturbations, and how compatible these behaviors are with model (e.g., HMM) assumptions, the selection of processing methods often involves substantial manual trace inspection as well as trial-and-error, reducing efficiency and possibly missing significant discoveries. To systematize this process of discovery, we propose the LSE metric, which evaluates the level of rarity of trace behaviors across multiple datasets (see Methods). Trace behaviors common across a group of datasets possess a high LSE, while those found only in a minority of datasets have a low LSE. Thus, clusters of traces with low LSE (e.g., on a UMAP projection) may signal the presence of behaviors strongly dependent on experimental conditions. To evaluate the utility of this metric on experimental data, we calculated the LSE for 9 experiments from a titration of KCl into the paused transcriptional elongation complex (D4 in Fig. 3a). In a common UMAP projection space, non-curated traces with low LSE in 50 mM KCl (Fig. 5d) are clustered in a different region than at 1,000 mM KCl (Fig. 5e), indicating KCl-dependent behaviors. These differences are highlighted even more strongly if only the 10% of traces with lowest LSE across all conditions are plotted (Supplementary Fig. 6e, f). Inspection of these low-entropy traces reveals a shift from predominantly low-FRET to high-FRET behaviors as KCl is increased, which is also reflected in FRET histograms of the two conditions (Supplementary Fig. 6d, h). However, unlike FRET histograms, the LSE-coded UMAP plots preserve individual trace information, can encapsulate a broader set of characteristics such as kinetics, and do not require prior trace curation (Supplementary Fig. 6a, b, c, d). The FRET histogram of the 10% lowest LSE traces clearly unveiled that the inflection point from low-to high-FRET resides at 400 mM KCl (Supplementary Fig. 6h), while the change from 300 mM to 400 mM KCl is recognizable, but not obvious in the corresponding traditional FRET histograms (Supplementary Fig. 6i). The LSE metric thus facilitates rapid identification of condition-specific trace behaviors without prior assumptions of which traits to consider, resulting in better informed analysis method selection.

### META-SiM Projector and LSE enable efficient discovery of a new intermediate in a complex smFRET system

Single-molecule data are inherently complex, but become even more so in the case of experimental systems that strive to reconstitute multistep biological processes such as translation or splicing. In such systems, observed behaviors result from the often complicated, dynamic interactions between one or more substrates and potentially several catalytic subunits and cofactors. Even with carefully designed controls to stall such pathways at specific steps, finite efficiency and a lack of synchronization dramatically increase the degree of heterogeneity of the observed single-molecule behaviors. As a result, even the intermediates of greatest biological insight, such as those appearing only in complexes stalled by the omission of a specific cofactor, may comprise only a small minority of the traces observed. A general method to quickly identify such condition-specific behaviors would greatly accelerate the pace of discovery. While the single-molecule clustering analysis (SiMCAn) method^6^, developed to systematically cluster related behaviors from highly heterogeneous datasets from a smFRET study of pre-mRNA splicing, represents progress towards this goal, it still relies on explicit modeling (e.g., HMM) and lacks a quantitative metric like LSE to quickly visualize and identify condition-specific behaviors.

To test its ability to accelerate discovery from such complex biological systems, we used META-SiM to analyze the smFRET data from the SiMCAn study of the yeast spliceosome, in which dynamic rearrangements of a pre-mRNA substrate were monitored as a function of progression through the splicing pathway^6^. In this study, a Ubc4 pre-mRNA substrate was labeled with a FRET pair comprising a Cy5 seven nucleotides upstream of the 5′ splice site (5′SS) and a Cy3 six nucleotides downstream of the branch point (BP; Fig. 6a), and immobilized *via* the biotin-streptavidin interaction on a quartz slide for smFRET imaging. The splicing pathway was stalled at specific steps by mutation of the pre-mRNA or of cofactors, or by depletion of specific cofactors from the extract, and the impacts of these perturbations on pre-mRNA dynamics monitored by smFRET (Fig. 6b). We used META-SiM to convert the smFRET traces from all experimental conditions to embeddings and calculated the LSE (Supplementary Fig. 7). Focusing on four of the most critical conditions related to the first and second steps of splicing (Fig. 6c), UMAP projections of the trace embeddings into either the smFRET Atlas (Fig. 6c) or system-specific (Fig. 6d) coordinates clustered low-entropy traces in specific regions. By contrast, for three of the four conditions, any differences in the FRET histograms (Fig. 6e) and TODPs (Fig. 6f) are subtle. By plotting only the 10% lowest-LSE traces across all datasets, we were able to isolate condition-specific behaviors in the UMAP projections (Fig. 6g-h), FRET histograms (Fig. 6i), and TODPs (Fig. 6j). Not only is the transition from a predominantly low-FRET state in the ΔPrp2(WT) condition to a predominantly high-FRET state in Prp16DN(WT) more prominent among the low-entropy traces, but there is evidence of a re-emergence of a minority low-FRET state in the WT(3’SS) and WT(WT) conditions (Fig. 6i). These observations suggest that the pre-mRNA is locked in a high-FRET conformation in the absence of a functional Prp2 helicase (Prp16DN(WT)), but can enter a state with a more extended (low-FRET) pre-mRNA in the presence of a functional Prp2 (WT(3’SS) and WT(WT) conditions; Fig. 6b red dashed box). This minority low-FRET population, not detectable in the original SiMCAn splicing study^6^, is consistent with observations from a later study that biochemically isolated an inactivated complex with an extended pre-mRNA^36^, and represents a rare and transient intermediate conformation involved in repeated 3’SS sampling that appears only in the presence of functional Prp2 and Prp22 helicases (Fig. 6b).

**Fig. 6.**
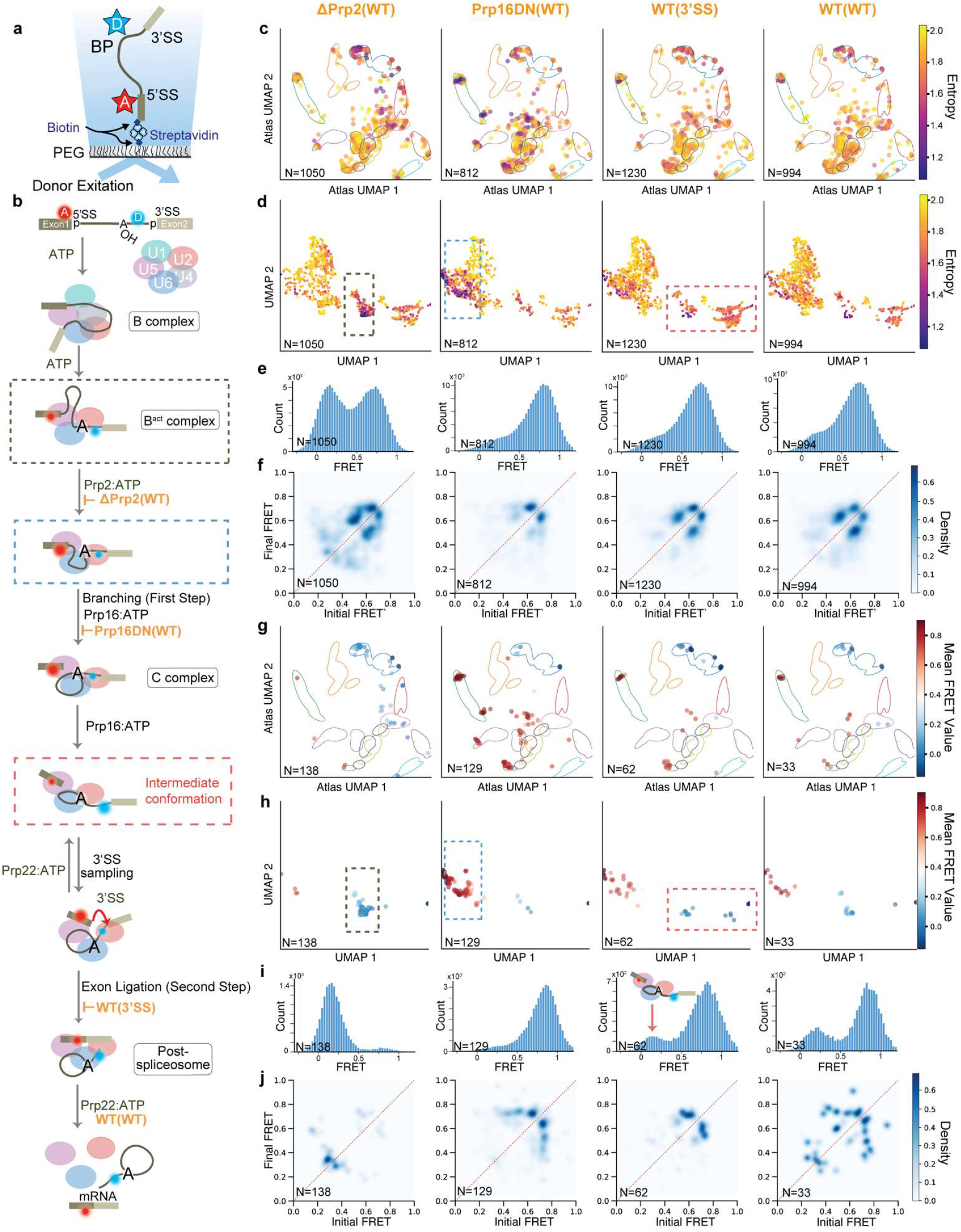
META-SiM as a tool for biological discovery in complex smFRET data. **a**, Labeling and immobilization scheme for use in a smFRET study of pre-mRNA splicing^6^. D = FRET donor, A = FRET acceptor, SS = splice site. **b**, Model of the multitude of pre-mRNA conformational changes and defined complexes during splicing, involving the first and second steps of catalysis. The dashed red box indicates a conformational intermediate discovered here in the experimental smFRET dataset. **c**,**d**, 2D UMAP projections of the manually curated traces from four critical conditions related to the first and second steps of splicing, where the traces are projected into smFRET Atlas (**c**) or system-specific (**d**) coordinates. **e**,**f** FRET histograms (**e**) and TODP plots (**f**) of the traces in (**c**) and (**d**). **g**,**h** 2D UMAP projection of the 10% of traces with the lowest LSE, projected into the smFRET Atlas (**g**) or system-specific (**h**) coordinates. **i**,**j** FRET histograms (**i**) and TODP plots (**j**) of the traces in (**g**) and (**h**).

Notably, the 10% lowest LSE traces are distributed differently across experimental conditions (Supplementary Fig. 8), which can help identify which conditions may yield the most unique or interesting behaviors. Our unprecedented discoveries from existing smFRET data were not evident until the traces with lowest LSE (most condition-specific behavior) were isolated and their properties examined, thus demonstrating the potential of the LSE metric and META-SiM Projector to readily identify rare, but biologically important behaviors in complex datasets.

## Discussion

Despite the power of single-molecule methods to deliver unique insights into the dynamic and heterogeneous properties of many biological systems^19,20,42,43^, their utility as a discovery tool has been hampered by data complexities that necessitate inefficient manual and *ad hoc* approaches for analysis. While automated tools and software have been developed to streamline or systematize data analysis, they have to date focused only on narrow sets of tasks^30,31^ or required the use of explicit physical or statistical models^26–29^ that, depending on the validity of model assumptions and the quality of fitting, may not accurately capture the full range of behaviors within a dataset. In addition, none of the existing tools provide a means of comprehensively visualizing the structure of SMFM datasets according to trace characteristics, or systematically identifying rare or condition-specific behaviors, without prior assumptions regarding what traits to consider.

Due to its multitask training, META-SiM not only excels at a variety of common single-molecule analysis tasks with little or no fine-tuning data but, by virtue of its high-dimensional output embeddings that encapsulate a rich array of information about each trace, establishes a new paradigm for systematic, open-ended biological discovery in single-molecule datasets. In particular, the META-SiM Projector enables the intuitive visualization of entire datasets and interactive inspection of related behaviors, while the LSE metric permits isolation of condition-specific trace behaviors—even if rare—for rapid hypothesis generation and further analysis. These capabilities not only enable more efficient discovery in conventional single-molecule studies, but may also lay the foundation for much higher-throughput discovery and screening efforts involving single-molecule observation, as well as facile data comparison across research groups.

Strikingly, this systematization allowed non-experts with no prior knowledge of splicing or the smFRET system to quickly identify a new intermediate that was not observed in the original study producing the data, despite the use of advanced data clustering methods^6^. The dataset visualization and LSE-based filtering tools, together with the automated trace classification and segmentation capabilities of META-SiM, render interpretation of single-molecule results much more accessible to researchers without expertise in single-molecule methods. The SCS metric further promotes accessibility and quality control by providing feedback on the consistency of trace classification and segmentation, while the web-based META-SiM Projector provides intuitive means of labeling datasets for model fine-tuning as well as sharing datasets and analyses with other researchers. Thus, while some domain expertise will still be required for full interpretation of results, META-SiM and its Projector have the potential to greatly streamline the analysis and discovery processes, which may contribute to the democratization of SMFM techniques by rendering their analysis more accessible to researchers outside of single-molecule biophysics.

Finally, the smFRET Atlas (Fig. 4m) offers a powerful means of visualizing and comparing entire single-molecule FRET datasets in a common space that reflects differences in FRET efficiency, number of states, kinetics, and other parameters relevant to smFRET analysis. Importantly, this and other visualizations in the META-SiM Projector can be performed without any prior curation, manual inspection, or model-based processing of traces, thus providing a rapid overview of even complex datasets prior to detailed analysis.

Foundation models have ushered in a revolution in natural language processing and other domains due to their versatility and consideration of a wide range of relevant information. In line with these precedents, we hope META-SiM will greatly increase the pace and scope of discovery in single-molecule biology. In the future, the model may be extended to an even wider range of applications in single-molecule biophysics. Furthermore, connecting the high-dimensional embeddings generated by META-SiM to large language models may allow users to summarize and interact with these complex datasets using natural language for even greater accessibility.

## Methods and Materials

### Network Formulation

Prior to feeding it into the network, each trace is divided into 50-frame patches and tokenized with a linear projection operation; each token thus reflects more information than a single frame, including short-range intensity changes and fluctuations. To encode the time dimension, we built a tokenizer similar to the one used for images in a vision transformer^44^: that is, we added a learnable position embedding^45^ to the tokens before the first layer of the attention model. In addition, we prepended a special learnable “trace-token” to the sequence of trace tokens to represent the intention to seek trace-level information, analogous to the class token in the language model BERT^45^. The corresponding output of the network from this trace-token is used for trace-level tasks, while output of other tokens was used for frame-level tasks. The attention block consists of multiple identical attention layers and maps the sequence of tokens to a sequence of embedding vectors. As a token represents a patch comprising multiple (50) frames, we further map the embedding vector of each patch to a list of embedding vectors for all frames with a linear projector to yield frame-level resolution for each trace. Both the trace-level and the frame-level embeddings are shared for all tasks.

We represent each time trace as a ℝ^*T*×*C*^ tensor where *T* is the total number of time steps and *C* is the number of color channels. *C* = 2 for two-color smFRET time traces and *C* = 1 for single color time traces. For ease of network training, we always use *C* = 2 and fill any non-existing color channel with zeros. We set up the tokenizer as a matrix multiplication operator over a segment of time steps. Each segment is a ℝ^*WC*^ vector denoted as *x*^segment^, where *W* is the width of the tokenizer, and the tokenization operation is:

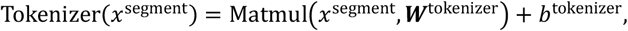

where ***W***^tokenizer^ ∈ ℝ^*WC*×*D*^ and *b*^tokenizer^ ∈ ℝ^*D*^ are the weights and bias of the tokenizer, respectively, *D* is the dimension of the embedding vectors, and Matmul denotes the matrix multiplication operation.

The tokenized sequence has a length of *T* / *W*. We prepend a learnable class token and add a learnable absolute position embedding to the sequence. Therefore, the input sequence to the attention model is of length *T′* = *T*/*W* + 1. We denote it as *x*^attention^ ∈ ℝ^*T′*×*D*^.

We use N layers of the standard multi-head self-attention model with layer normalization to operate on *x*^attention^:

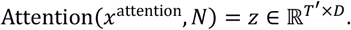

We use the first step of *z* as the trace-level embedding denoted as *z*_*o*_, and denote the remaining *T*^*′*^ − 1 time steps as *z*_1_, …, *z*_*n*_. We further implement an unstack operator, the reverse operator of the tokenizer, to convert *z*_*i*_ into the embedding for each time step in the original time trace. The unstack operator reads:

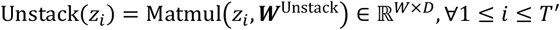

where ***W***^Unstack^ ∈ ℝ^*D*×*W*×*D*^ is the weight.

### Synthetic Training Data Generation

To obtain enough training data for all tasks, we used a trace simulator^31^ to generate over 1 million synthetic traces. Ground truth labels were generated from simulation parameters or ideal measurements with no noise. The simulation is based on the stochastic Markov chain process describing the state transitions of a time trace. To sample simulation parameters, we first sample the number of distinct states uniformly ranging from 1 to 4, and then the initial state probability distribution *p*_init_(*s*) and transition probability matrix *p*_transfer_(*s*_1_, *s*_2_) from independent uniform distributions, where *s* is the initial state and *s*_1_ and *s*_2_ are the states before and after a transition. We also sample the total intensity *I*_max_ from a uniform distribution. We can thus generate a two-color time trace *I*_donor_(*t*), *I*_acceptor_(*t*) or one-color time trace *I*(*t*) that has states and transitions but no noise or photophysical behavior.

Next, we simulated the photophysical behavior including photobleaching, blinking and quantum yield. We sampled the photobleaching lifetime for each color channel from independent exponential distributions with expected lifetime *τ*_bleach_ sampled from a uniform distribution ranging from 1 frame to 2000 frames. Then we sampled the blinking of Cy3 dye by sampling the non-blinking lifetime and the blinking lifetime from independent exponential distributions with expected lifetime *τ*_non-cy3-blink_ and *τ*_cy3-blink_ sampled from uniform distributions. Finally, we drew the relative acceptor brightness parameter from a uniform distribution from 0.9 to 1.0 and multiplied this value with the original *I*_acceptor_ to generate the new *I*_acceptor_.

Lastly, we added Gaussian noise to the intensity traces, as well as a constant offset value, sampled from a uniform distribution yielding 1.8 < SNR < 6, to represent background signal.

### Multitask Training and Hyperparameter Optimization

We used the synthetic training data to create various learning tasks (Supplement Table 1) for the network to recognize important features of single molecule time traces. These learning tasks include trace-level tasks such as recognizing kinetic states, SNR, photobleaching lifetime and blinking, and frame-level tasks such as recognizing regions of active kinetic transitions and predicting idealized FRET states. We designed a multitask pre-training strategy to optimize the training on all trace- and frame-level tasks simultaneously using the combined loss of all tasks. One common challenge when combining the loss of multiple tasks is that one or a few tasks can dominate the total loss, leading to the network being poorly optimized on other tasks. For example, the labels measuring photobleaching lifetime have a range of [1, 2000] frames, while the labels measuring FRET ratio have a very different range of [0, 1]. If we use a mean-absolute-error loss, the network will likely be optimized for photobleaching lifetime but poorly optimized for FRET value prediction. In addition, the training dataset includes both classification and regression tasks, necessitating a unified loss function. To address this concern and unify the loss function across all tasks, we employed a cross-entropy loss function and quantized all tasks as multi-class classification problems, unifying classification and regression tasks. Specifically, we quantized the labels for all tasks and used a KL divergence loss function for all the tasks. The loss function, which is denoted as:

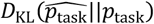

and measures the difference between the training labels’ empirical probability distributions 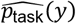 and the model’s predictions *p*_task_(*y*; *x*, ***θ***), is similarly valued for all tasks even though their labels have divergent values. Putting it together, the combined loss function to optimize, ignoring the constant terms, is:

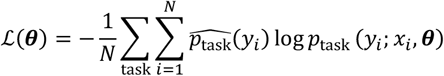

as a function of network parameter ***θ***.

To generate *p*_task_(*y*; *x*, ***θ***), we used an independent linear operator for each task followed by a softmax operation *σ*(⋅) to generate a probability distribution. We denote the trace- or frame-level encoding outlined in Network Formulation as METASIM(*x*_i_), the predicted probability distribution is:

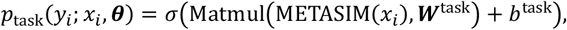

where ***W***^task^ and *b*^task^ are the task-specific weight and bias.

Another common challenge in multitask learning is competing tasks leading to poor optimization. To avoid training tasks in the next iteration of network training unintendedly competing with those of the previous iteration, we made sure each iteration’s training data were sampled evenly from all training tasks (Fig 2b). The network was optimized using the ADAM optimizer^46^ with *β* = 0.9 and *β* = 0.999. The network was trained on 1.5 million synthetic time traces, equivalent to 55 gigabytes of float32 data, for 2 days on a single NVIDIA A100 GPU.

The training and validation loss decay approximately linearly as a function of the logarithm of training steps, showing the slow-down of network optimization as the compute time increases (Fig. 2c). We optimized the important hyper-parameters of the network by a grid search of the depth, width, the number of attention heads, the activation function in each attention layer, and the width of the tokenizer (Supplementary Fig. 9a, b, c, d). The lowest validation loss, indicating the best performance, was obtained for a self-attention network 4 layers deep and 96 dimensions wide with 4 attention heads, with a Gaussian Error Linear Units activation function, comprising a total of 7 million parameters. Further increasing the number of network parameters by increasing the depth, the width, or the number of attention heads did not additionally decrease the validation loss (Supplementary Fig. 9e), a sign that the network’s learning capacity is high enough to learn the features of the time traces in the training data. In other words, we do not expect that a larger foundation model would dramatically increase the performance on common single molecule data analysis tasks unless more complex synthetic data generation models—resembling more complex experimental systems—must be included for model training.

### Supervised fine-tuning

To fine-tune META-SiM for common downstream tasks (Fig. 3), we assigned a new head to the model containing one linear classification layer for each task, and performed a logistic regression to the head with a small amount of labeled data on the order of 100-1,000 traces. Specifically, we perform supervised fine-tuning with frozen encoders, a widely used fine-tuning strategy for small datasets. In this fine-tuning, referred to as “logistic regression” (LR), only ***W***^task^ and *b*^task^ in the last classification layer are varied during training, while the remaining network parameters in METASIM(⋅) are kept fixed. This vastly reduces the amount of data and computational power required for training; for example, fine-tuning using 100-1,000 labeled traces can be completed on a typical laptop in a few minutes. While the network also supports fine-tuning that updates *all* the model’s parameters, this is not anticipated to be a common need, and is not used for any of the examples in this study.

### Evaluation of Performance in Downstream Tasks

To evaluate the performance of META-SiM fine-tuned for trace classification and segmentation—e.g., to identify traces and segments thereof that provide meaningful information, an essential early step in analysis of smFRET traces—we employed five experimental smFRET datasets (Fig. 3a). For benchmarking, we used the previously reported task-specific deep learning model DeepFRET^31^, but initially found that its direct application to D1-D5 yielded low concordance (below 80%, Fig. 3b). This suggests that DeepFRET may not generalize well to these five experimental systems without further fine-tuning. Therefore, for a fair comparison we performed the same logistic regression operation with the same fine-tuning dataset as META-SiM to yield a fine-tuned DeepFRET (Fig. 3b, c). Across the five datasets, META-SiM_LR_CS and DeepFRET achieved similar concordance (Fig. 3b), FRET-distribution overlap (Fig. 3c), and area under the receiver-operating characteristic curve (ROC AUC) (Supplementary Fig. 2a), indicating comparable performance of META-SiM to a state-of-the-art task-specific network when both models are similarly fine-tuned for the task. Compared to our previous network Auto-SiM^30^, a task-specific deep learning network for classification and segmentation that used more than 50,000 manually labeled experimental traces for training, META-SiM_LR_CS achieved a similar ROC AUC while needing only 82-215 (Supplementary Table 2) labeled traces for fine-tuning and no experimental data in pre-training. This >200-fold reduction in the amount of manually labeled data required for fine-tuning makes META-SiM much more practical as a customizable labor-saving tool than task-specific models, while retaining high performance. Note that the web app META-SiM Projector streamlines the creation of these manual labels for new datasets in case any fine-tuning is needed.

To evaluate the performance of META-SiM in trace idealization without fine-tuning, we used META-SiM directly to idealize 19 traces from a dataset in a public benchmark study^14^ of a two-state smFRET experimental system. Rate constants of FRET transitions were obtained by exponential fitting of the cumulative dwell time distributions in the low- and high-FRET states from these idealizations (Supplementary Fig. 1c). As ground truth values of the rate constants and FRET states in the experimental dataset are unknown, the results from META-SiM were compared with those from 14 other methods (Fig. 3f, g), following the assessment conventions in the prior benchmarking study^14^. The kinetic constants derived from the idealized FRET states by META-SiM deviate a modest 3.6% and 6.8% from the mean predicted for the kinetic constants k_12_ and k_21_, respectively, while the FRET state values predicted by our model deviated only 0.86% and 0.24% from the mean predictions, respectively. By comparison, the mean deviation from the mean prediction among the 14 other tools were 9.2% and 12.1% for kinetic constants k_12_ and k_21_ and 1.4% and a 0.42% for the FRET state values, respectively. The fine-tuned version (META-SiM-FT) for trace idealization gave similar results compared to the non-fine-tuned one (Supplementary Fig. 1g, h). These findings demonstrate that META-SiM performs on par, if not better, than all other currently available, state-of-the-art single molecule analysis tools.

To evaluate the performance of META-SiM in counting photobleaching steps—apparent as instantaneous drops in fluorescence corresponding to the bleaching of single fluorophores—we compared the predictions of the model to those of manual counting by researchers. This strategy was implemented because incomplete labeling and activity of fluorophores makes it challenging to ascertain a ground truth in these experiments. The fine-tuned META-SiM achieved a concordance of 68.0% for exact count matching and 90.0% for exact or off-by-one matching (Supplementary Fig. 1e). For reference, concordance between independent researchers was higher, at 84.5% for exact count matching and 95.0% for exact or off-by-one matching (Supplementary Fig. 1e). However, the histogram of photobleaching steps counted by META-SiM is in close agreement with the manual labeling (Fig. 3i), and the mean number of steps (3.9 +/- 0.14, 1 s.d.) aligns well with the mean value from manual labeling (3.9 +/- 0.13, 1 s.d.).

To evaluate the performance of META-SiM for single-molecule recognition through equilibrium Poisson sampling (SiMREPS)^38,47^, we tested its ability to distinguish traces resulting from *EGFR* fragments containing the cancer-related single-nucleotide substitution T790M (mutant) and those from wild-type *EGFR* fragments (Fig. 3j,k)^48^. Compared to conventional classification based on HMM fitting followed by kinetic thresholding, META-SiM consistently accepts more true positives and fewer false positives, corresponding to an ∼40% increase in sensitivity and an ∼3.75-fold increase in specificity (Fig. 3k). META-SiM also increases the sensitivity by 1.4-fold for a standard curve of T790M in the absence of wild-type sequence (Supplementary Fig. 1f). This high performance, despite fine-tuning only for the last layer of the network with less than 200 trainable parameters, suggests that the META-SiM foundation model can be easily adapted to a wide variety of specialized trace-classification tasks.

### META-SiM Projector

To help researchers easily visualize dimensionally reduced projections of their own single-molecule data and apply labels for fine-tuning of the foundation model to their own projects, we built an online interactive tool called META-SiM Projector (https://www.simol-projector.org). This tool allows users to visualize the embeddings generated by META-SiM from their own experimental data as 2-D or 3-D projections created with PCA, t-SNE or UMAP algorithms. In these projections, each point represents a single-molecule trace. When a point is selected, the corresponding trace is shown along with a list of other traces ordered by distance in embedding space from the selected trace, facilitating interactive browsing of the dataset by trace similarity. Users can also use a label creation tool to apply numerical, text, and/or Boolean labels to traces; upon so doing, label Self-Consistency Scores (SCS) are calculated in real time for feedback on the consistency of labeling. These labels can then be directly used for fine-tuning of META-SiM *via* the provided Google Colab notebooks (https://www.simol-projector.org), and can be saved with the raw time traces to a local hard drive. Even in cases where META-SiM performs well without fine-tuning, manual labeling of a small subset of the data (followed by complete labeling with the fine-tuned model) can be very useful as an annotation tool that facilitates recognition of patterns in a dataset from its projection plot (e.g., UMAP). PCA, t-SNE and UMAP embedding projections can be exported as images, and individual time trace plots can be saved as vector graphics files. META-SiM Projector also allows isolation of individual sub-populations within a dataset and re-calculation of the embedding projection to investigate variation within the sub-population. Finally, when a user analyzes data with this tool, a unique link can be generated for easy sharing of traces and embedding projections between users and research groups. More comprehensive guidelines and details about software functions are described in the META-SiM Projector Quick User Guide on the website.

### Principal Projection of Embedding Vectors

To provide more direct control over the projection of information contained in the embedding vectors generated by the network into a lower-dimensional space, we developed a method to find linear operations 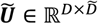 that project a D-dimensional META-SiM trace-level embedding vector ***z*** = METASIM(*x*) ∈ ℝ^*D*^ into a lower-dimensional embedding vector. The projection operator for a group of tasks {task_*i*_}, *i* ∈ [1, *n*] is obtained by finding the principal directions that contribute the most to these tasks. This is achieved by computing a single-value decomposition of the concatenated task weight matrix 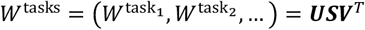, where the weight matrices are obtained from pre-training.

To reduce the rank of the projection matrix, we keep the largest diagonal components of *S* that account for *λ* = 95% of total second 2^nd^ momentum, i.e., 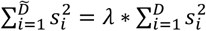 where *S* is in decreasing order. The matrix ***Ũ*** is obtained by using the corresponding columns of the left unitary matrix ***U*** of the singular-value decomposition. The tasks are grouped together based on their common physical meanings for time traces, such as kinetic rate constant, FRET value distributions, signal-to-noise ratio and photophysical properties (Supplementary Table 1).

To measure the alignment of two low-rank projection matrices ***Ũ***_1_ and ***Ũ***_2_, we use the 2-norm of their product matrix, i.e., 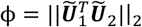 where we denote the alignment as ϕ. For orthogonal ***Ũ***_1_ and ***Ũ***_2_, *ϕ* = 0. For equivalent ***Ũ***_1_ and ***Ũ***_2_ up to a unitary rotation, *ϕ* = 1. For general ***Ũ***_1_ and ***Ũ***_2_, 0 ≤ *ϕ* ≤ 1.

### Label Self-Consistency Score (SCS)

The label SCS is defined as the probability of observing the same label between two traces if they are *k-*nearest-neighbors in the embedding space, i.e., one trace is among the k nearest neighbors of the other based on the k-nearest neighbors algorithm (k-NN)^49^, averaged across all traces in the dataset. Given a dataset of traces {*x*_*i*_} and their labels {*y*_*i*_} where *i* ∈ [1, *N*], we first run network inference to obtain the trace-level embedding vectors *z*_*i*_ = METASIM(*x*_*i*_) for each trace. For each *z*_*i*_, we find the *k* nearest neighbours NN(*z*_*i*_, *k*). The label SCS for a label *l* is defined as:

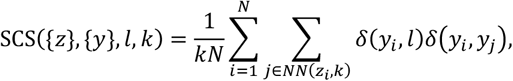

where *δ*(*i, j*) is the Kronecker delta function. We used *k* = 1 throughout this study.

### Local Shannon Entropy (LSE) of Single-Molecule Traces

Given a dataset of traces {*x*_*i*_} and their experimental conditions {*y*_*i*_}, where *y*_*i*_ is a categorical label belonging to a finite set of experimental conditions *C*, we obtain the trace-level embedding of each trace *z*_*i*_ = METASIM(*x*_*i*_). Then, for each *z*_*i*_, we find the *k* nearest neighbors NN_*i*_ = NN(*z*_*i*_, {*z*}, *k*). The LSE for a trace itself and its local nearest neighbors, following the standard definition of Shannon entropy, is written as follows:

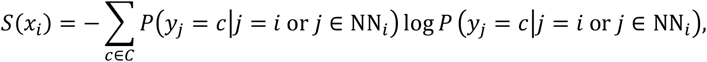

where the probability *P* can be calculated by:

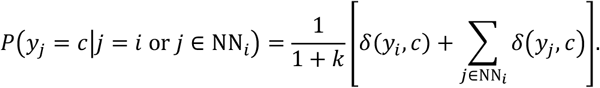

where *δ*(*i, j*) is the Kronecker delta function.

In this study, we used *k* = 50 for all Shannon entropy calculations. In principle, *k* should be much smaller than the total number of traces for locality. A very small *k*, however, should also be avoided to minimize fluctuations of the Shannon entropy value due to randomness in small samples. A rule-of-thumb for initial value of *k* is to use 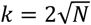 where *N* is the total number of traces in the dataset. Users can visualize the Shannon entropy values (as in Figs. 5-6) to empirically determine whether the selected *k* value can reliably detect heterogeneous behaviors in a given dataset.

### Single-molecule FRET Atlas

The smFRET Atlas allows sharing and comparison of diverse smFRET datasets in a common, global UMAP projection space. In this Atlas, traces with a given behavior pattern will always appear within certain regions of the coordinate space. To construct this Atlas, we generated 1 million synthetic smFRET traces with varying dynamics, signal-to-noise ratio (SNR) and FRET value distributions, aiming to sample as diverse a set of behaviors as possible to serve as a basis for the global UMAP projection. The UMAP model was optimized with unsupervised training using these synthetic traces.

To annotate this space for greater comprehensibility, the synthetic traces were grouped into 22 categories with a systematic nomenclature based on the simulation parameters used, “**number of states - SNR - FRET values – transition rate**”. While the number of states (1-3) is already an integer parameter, SNR, FRET values, and transition rates were divided into categories as follows. SNR was divided into “clean” (SNR ≥ 4) and “noisy” (SNR < 4); FRET value into “high” (value > 0.65), “medium”, (0.35 ≤ value ≤ 0.65) and “low” (value < 0.35); and transition rate into “fast” (average rate ≥ 0.05 frame^−1^) and “slow” (average rate < 0.05 frame^−1^). Categories are labeled using the first number or letter of each classification label. For example, “2-c-mh-f” means a trace with 2 kinetic states, clean SNR, one medium state and one high state, and a fast transition rate. A full list of category labels is presented in Supplementary Table 3. For annotation, we simulated 22,000 traces in total—1,000 for each category—and fit the probability density of the projected UMAP scatter plot with a two-dimensional 4-component Gaussian mixture model. We chose a small probability density *p* = 10^−2^ to draw the contour of the fitted probability density, which shows an approximate boundary for each category (Fig. 4m, Supplementary Fig. 5).

An end user can directly load the trained UMAP to project their own data and plot the fitted boundary to relate their own data to the categories used in the synthetic smFRET traces, increasing interpretability of this low-dimensional projection of their dataset(s).

### Experimental Datasets

Experimental datasets for downstream task analysis evaluation are from 9 different biological systems (Supplementary Table 2). For trace classification and segmentation, 5 systems were included for fine-tuning and testing: a toehold-exchange-based DNA walker^15^ (D1), a DNA swinging arm^50^ (D2), a preQ_1_ riboswitch^51^ (D3), a paused transcriptional elongation complex^35^ (D4), A Mn^2+^ riboswitch^52^ (D5). For stoichiometry analysis, we used an unpublished dataset (D6) from a 6-subunit protein complex bearing up to one HaloTag Alexa Fluor 660 (Alexa660) label per monomer. For kinetic fingerprinting, the dataset (D7) is from the detection of the EGFR point mutation T790M DNA sequence^48^. For trace idealization, the dataset (D8) is from a benchmark study of 14 tools where this specific testing dataset is from ACTR-NCBD Binding at 10-ms binning^14,53^. For the biological discovery assisted by LSE, the dataset is from an smFRET study of pre-mRNA splicing in yeast^6^.

## Data Availability

Experimental traces (stored in a customized data format) and META-SiM model weights (stored in HDF5 format) are available in our GitHub repository https://www.github.com/simol-lab/meta-sim.

## Code Availability

Code for generating embedding vectors and UMAP projections, applying META-SiM for downstream tasks and calculating metrics including the SCS and LSE and is available in our GitHub repository https://www.github.com/simol-lab/meta-sim.

## Acknowledgements

We thank Jingxuan Tang for stepwise photobleaching data. We thank Liuhan Dai and Pavel Banerjee for multiplexing SiMREPS data. We thank Adrien Chauvier for SiM-KARTS data. We thank Zhuoru Li for aesthetic consulting of main text figure 1 design. This work was supported by NIH grant R35 GM131922 to N.G.W.

## Author contributions

J.L. and L.Z. conceived the ideas and analyzed and interpreted the data. L.Z. and J.L. wrote Python programs for data processing, deep learning network training and single-molecule trace simulation. J.L., L.Z., A.J.B. and N.G.W. co-wrote the paper. All authors discussed the results and edited the manuscript.

## Additional information

Supplementary information is available in the online version of the paper. Reprints and permissions information is available online at www.nature.com/reprints. Publisher’s note: Springer Nature remains neutral with regard to jurisdictional claims in published maps and institutional affiliations. Correspondence and requests for materials should be addressed to J.L. and N.G.W.

