## SUPPLEMENTARY INFORMATION for "Foundation model for efficient biological discovery in single-molecule data"

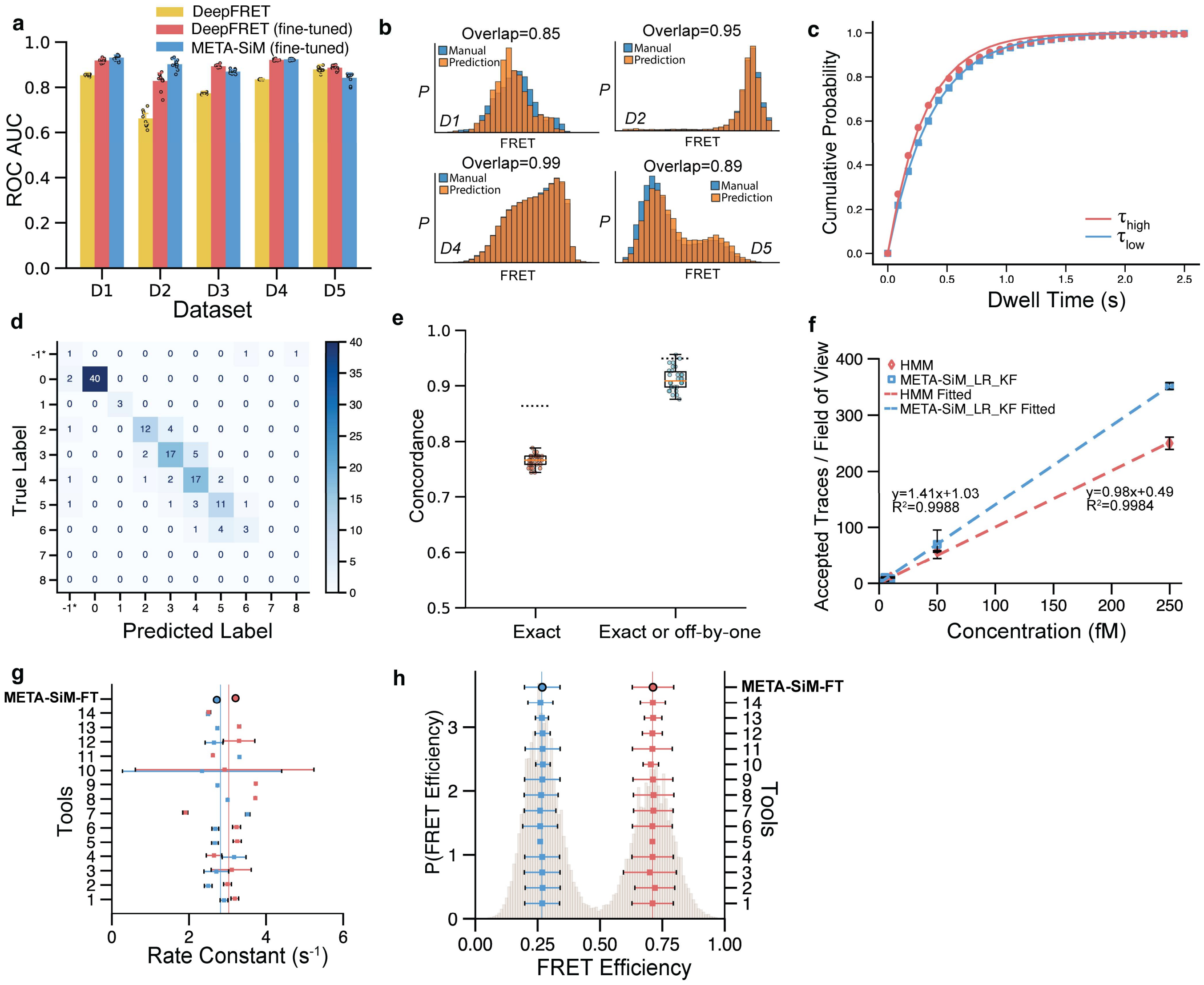


**Supplementary Fig. 1| Performance of META-SiM in various downstream tasks. a,** The area under the receiver-operating characteristic curve (ROC AUC) for trace classification by META-SiM and DeepFRET compared to manual analysis. **b,** Representative FRET histograms based on traces curated and segmented by META-SiM versus manual analysis. **c**, A distribution of dwell time predicted by the model is fit with single exponential distributions to yield transition rate constants. **d,** A representative confusion matrix comparing the labels from manual counting ("True Label") to the labels predicted by META-SiM. **e,** Concordance between manual counting and predictions by META-SiM that either match exactly or differ by no more than one step. **f**, Standard curve for T790M generated by META-SiM and HMM analysis. **g, h**, Evaluation of performance in trace idealization for META-SiM (Fine-Tuned), and benchmarking against 14 other common analysis tools[1], on the basis of measured rate constants (**g**) and FRET efficiencies (**h**): (1) Pomegranate; (2) Tracy(HMM); (3) FRETboard; (4) Hidden-Markury; (5) SMACKS(SS); (6) SMACKS; (7) Correlation; (8) Edge finding(CK); (9) Edge finding(k-means); (10) Step finding; (11) STaSI; (12) MASH-FRET(bootstrap); (13) MASH-FRET(prob); (14) postFRET.

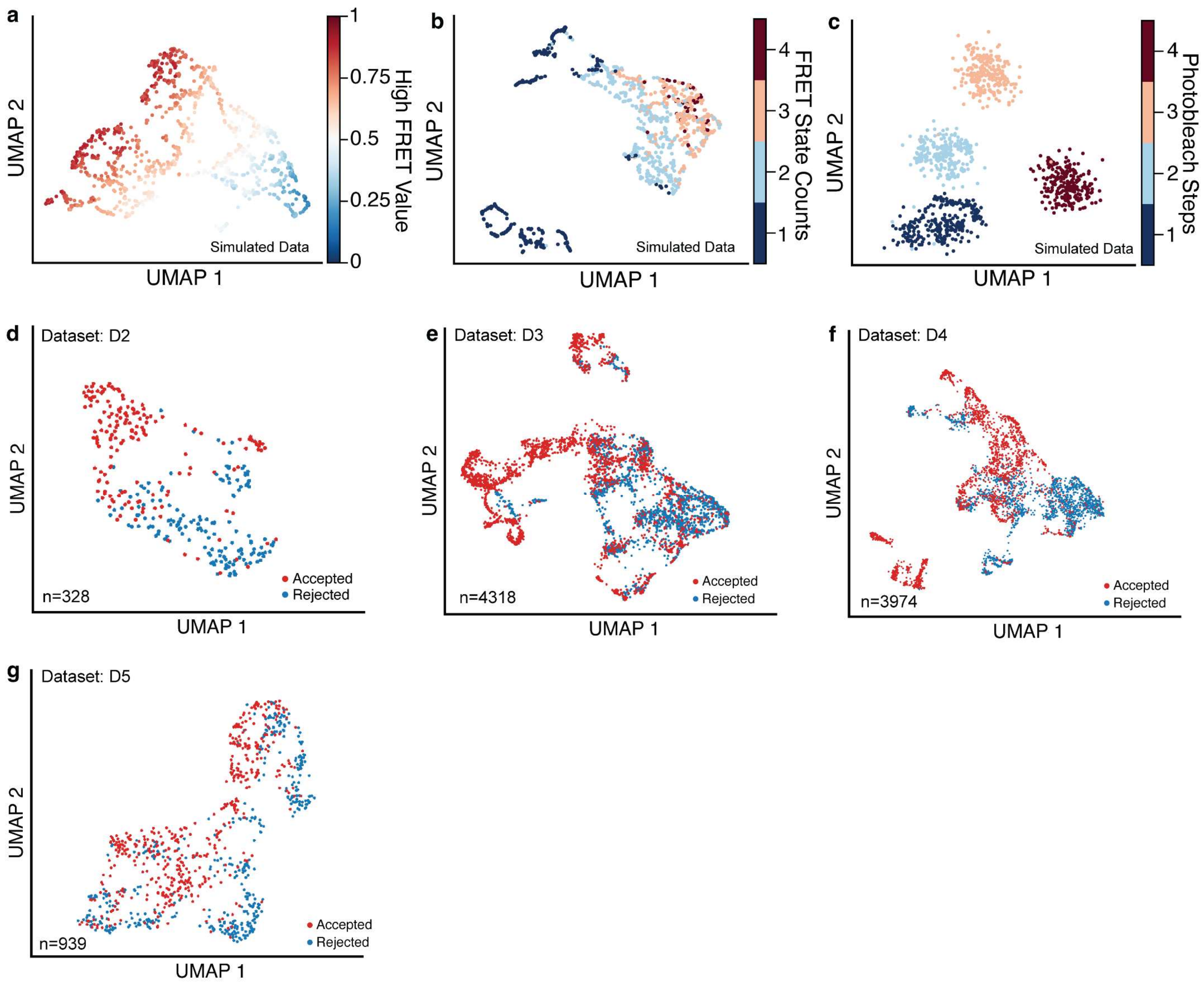


**Supplementary Fig. 2 | Evaluation of UMAP projection on both simulated and experimental data.** **a,b,c,** 2D UMAP projections of simulated traces with varying high-FRET value (**a**), number of FRET states (**b**), and number of photobleaching steps (**c**). **d,e,f,g,** 2D UMAP projection of traces from dataset D2[2], D3[3], D4[4], D5[5] that were manually accepted (red) or rejected (blue) for further analysis.

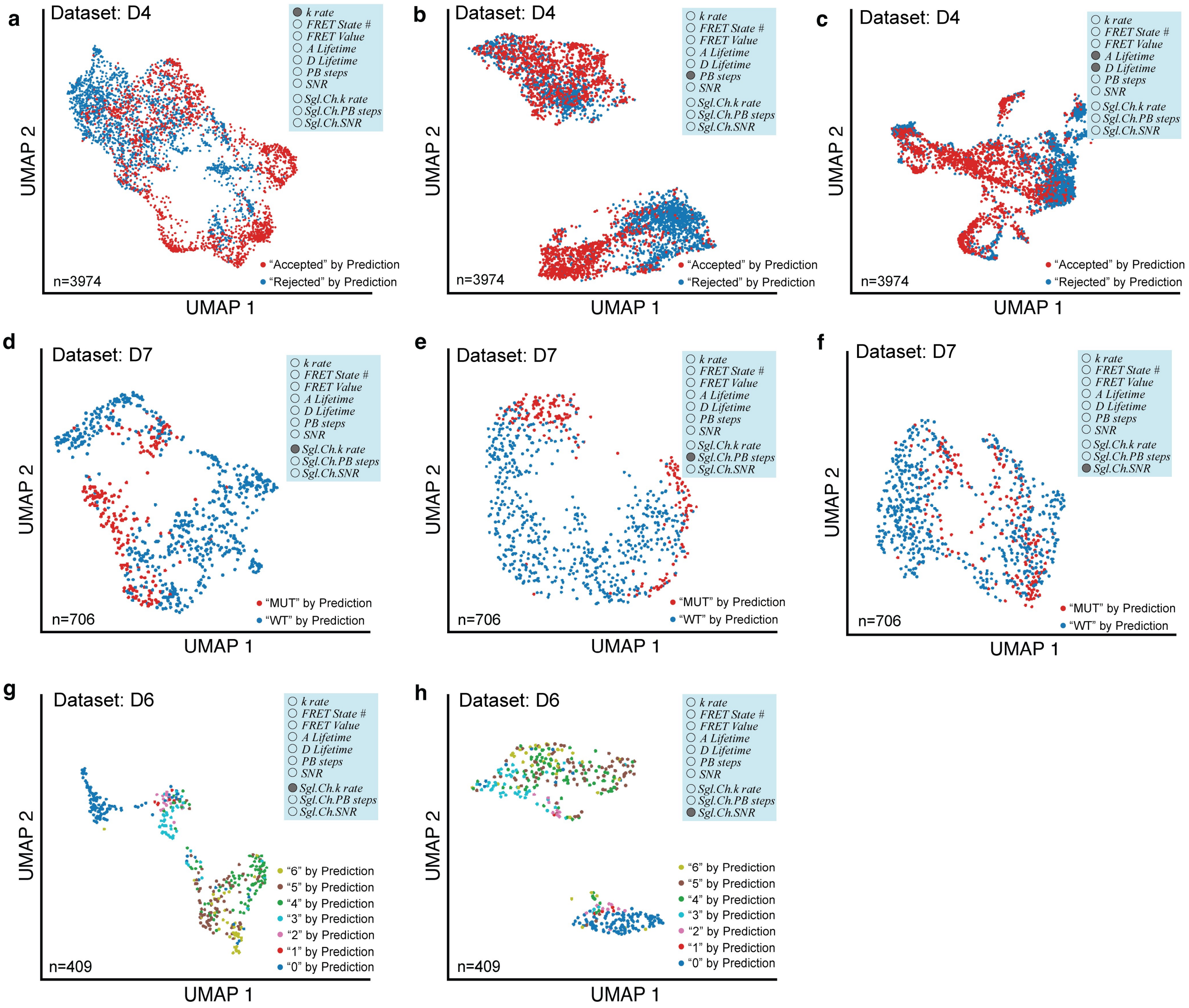


**Supplementary Fig. 3 | Varying principal projections from the same dataset. a,b,c,** 2D UMAP projection of dataset D4[4] based on different attributes: kinetic rate (**a**), photobleaching steps (**b**), donor and acceptor fluorophore lifetime prior to photobleaching (**c**). **d,e,f,** 2D UMAP projection of dataset D7[6] based on different attributes: single-channel kinetic rate (**d**), single-channel photobleaching steps (**e**), single-channel SNR (**f**). **g,h,** 2D UMAP projection of dataset D6 with different attributes: single-channel kinetic rate (**g**), single-channel SNR (**h**).

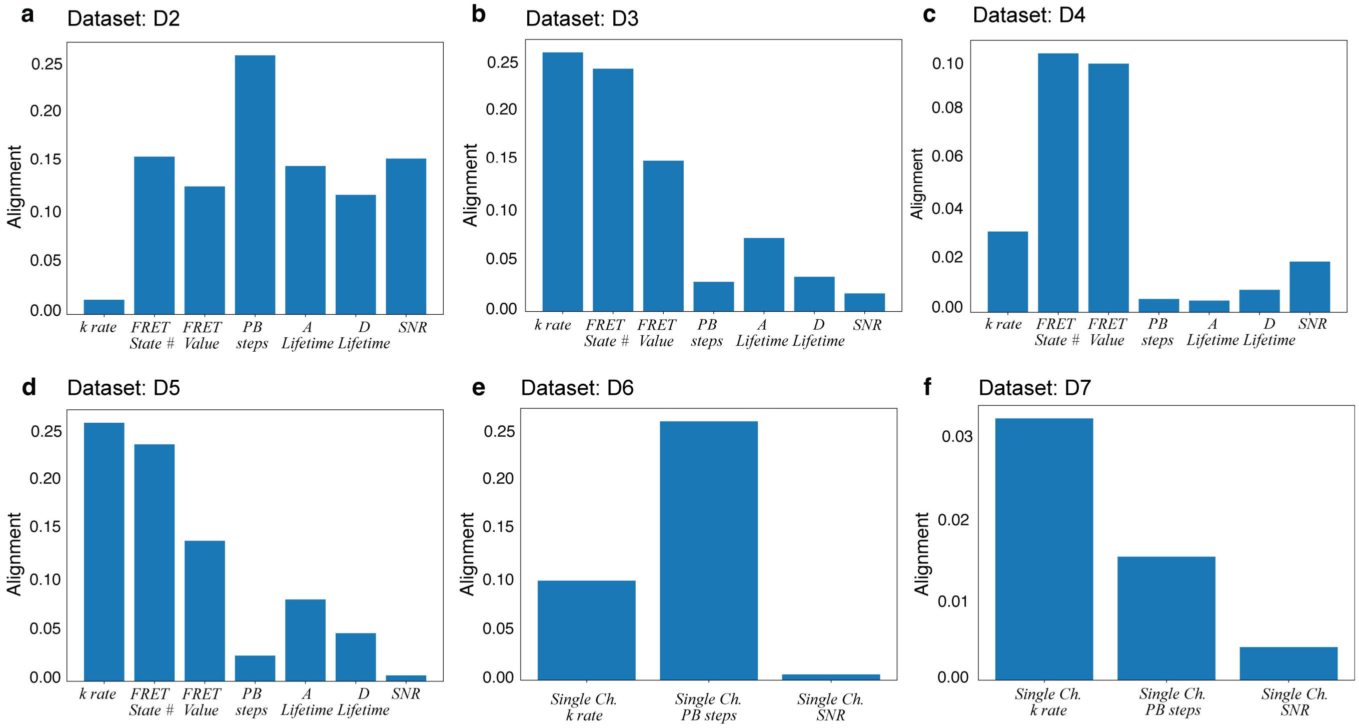


**Supplementary Fig. 4 | Quantitative assessment of the alignment of individual data attributes to network predictions. a, b, c, d, e, f,** Alignment of specific data analysis attributes with the META-SiM classifier fine-tuned for the different datasets D2, D3, D4, D5, D6, D7 to achieve the classification task.

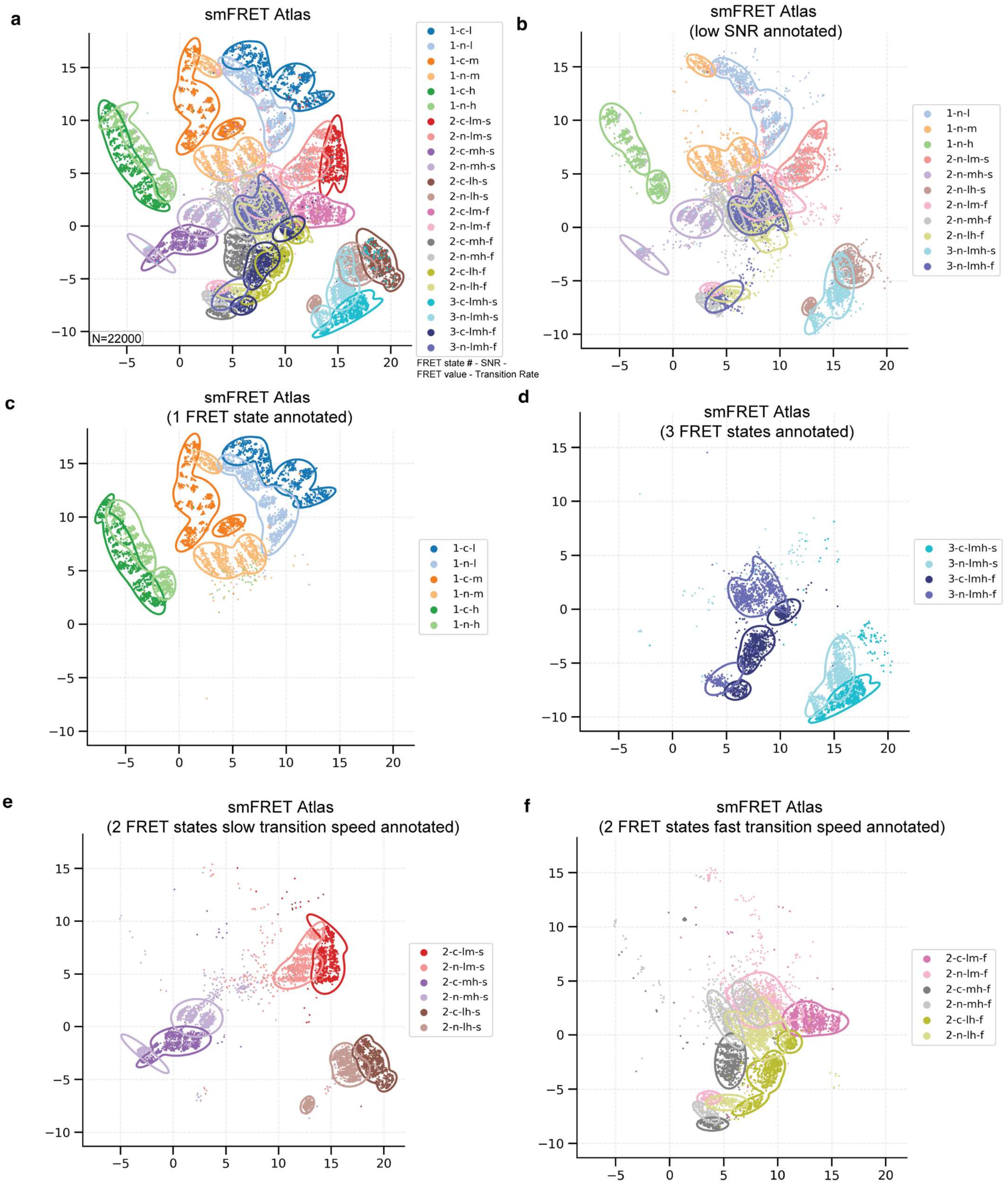


**Supplementary Fig. 5 | Different annotations of the smFRET Atlas. a,** An smFRET Atlas constructed with 22,000 traces derived from simulation. **b,c,d,e,f,** the same Atlas where only (**b**) low-SNR traces, (**c**) traces with a single FRET state, (**d**) traces with 3 FRET states, (**e**) traces with two FRET states and slow transitions, or (**f**) traces with two FRET states and fast transitions are plotted. Codes for cluster names (1-c-l, etc.) are listed in **Supplementary Table 3**.

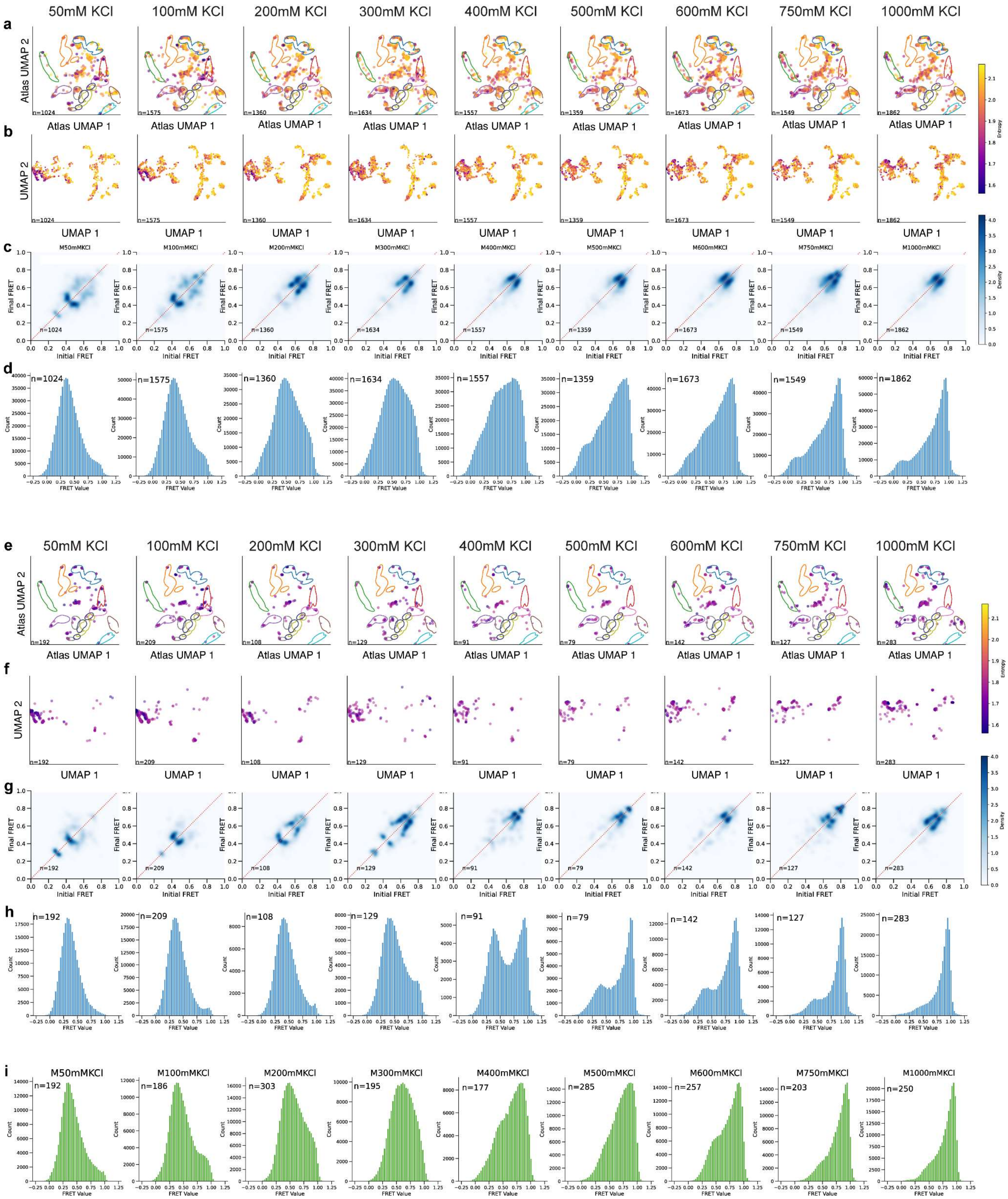

**Supplementary Fig. 6 | Titration of KCl into the paused transcriptional elongation complex system (D4). a,b,** 2D UMAP projections of embeddings from uncurated traces under different KCl concentrations in Atlas coordinates (**a**) and system-specific coordinates (**b**). **c,** TODPs of traces from the different KCl concentration conditions. **d,** FRET histogram of traces from the different KCl concentrations. **e,f,** 2D UMAP projections of the 10% of embeddings with lowest LSE from traces under the different KCl concentrations in Atlas coordinates (**e**) and system-specific coordinates (**f**). **g,** TODPs of 10% lowest LSE traces from the different KCl concentration conditions. **h,** FRET histograms of the 10% of traces with lowest LSE from the different KCl concentrations. **i,** FRET histograms of manually curated traces from the different KCl concentrations.

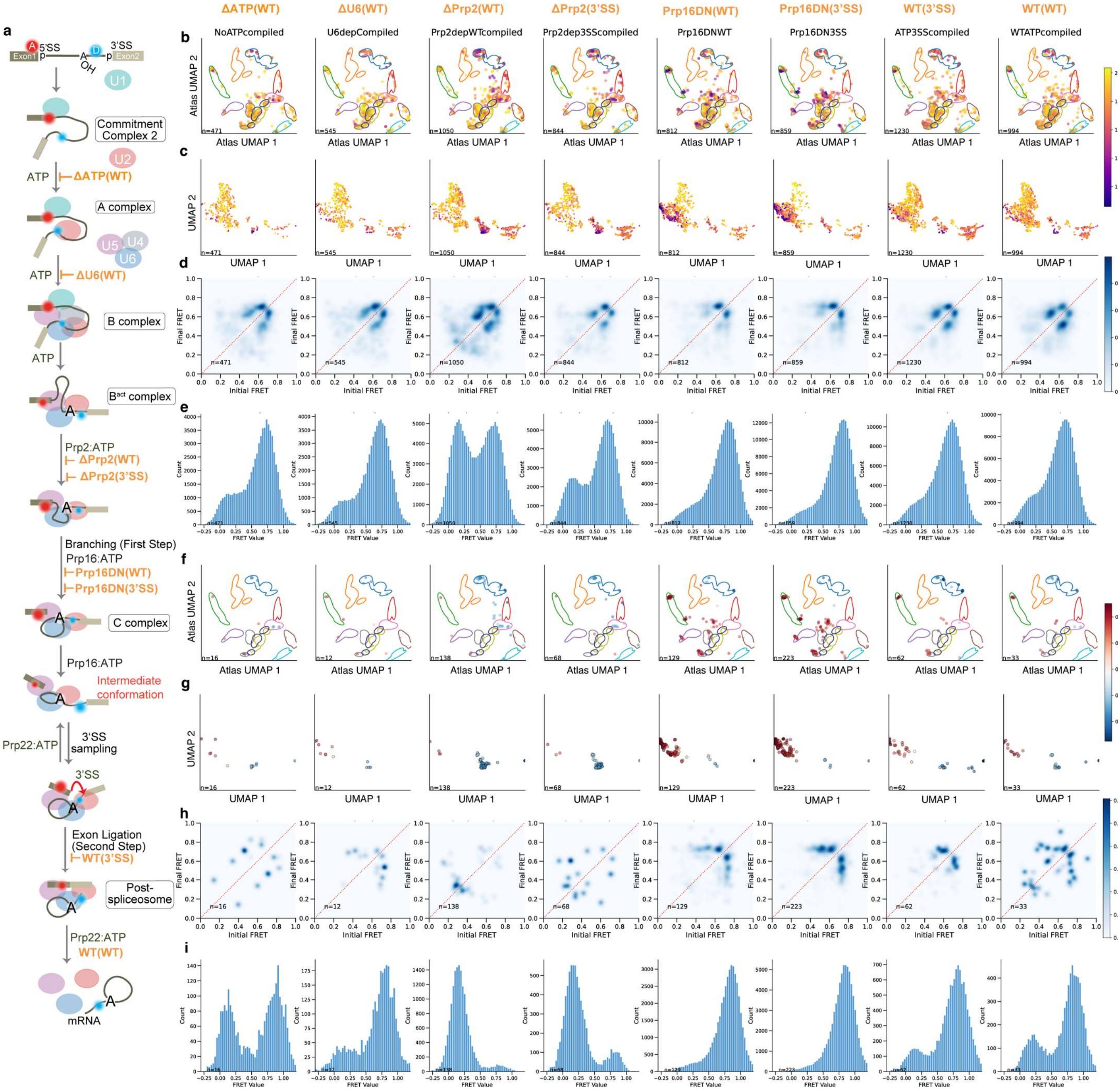


**Supplementary Fig. 7 | Full smFRET characterization of the yeast pre-mRNA splicing pathway. a,** Diagram of the splicing pathway. Experimental conditions used to block any further progress beyond specific steps in the pathway are annotated in orange font[7]. **b, c,** 2D UMAP projections of trace embeddings from the different experimental conditions in Atlas coordinates (**b**) and system-specific coordinates (**c**). **d,** TODPs of traces from the different experimental conditions. **e,** FRET histograms of traces from the different experimental conditions. **f, g,** 2D UMAP projections of the 10% of traces with lowest LSE under the different experimental conditions in Atlas coordinates (**f**) and system-specific coordinates (**g**). **h,** TODPs of the 10% of traces with lowest LSE from the different experimental conditions. **i,** FRET histogram of the 10% of traces with lowest LSE from the different experimental conditions.

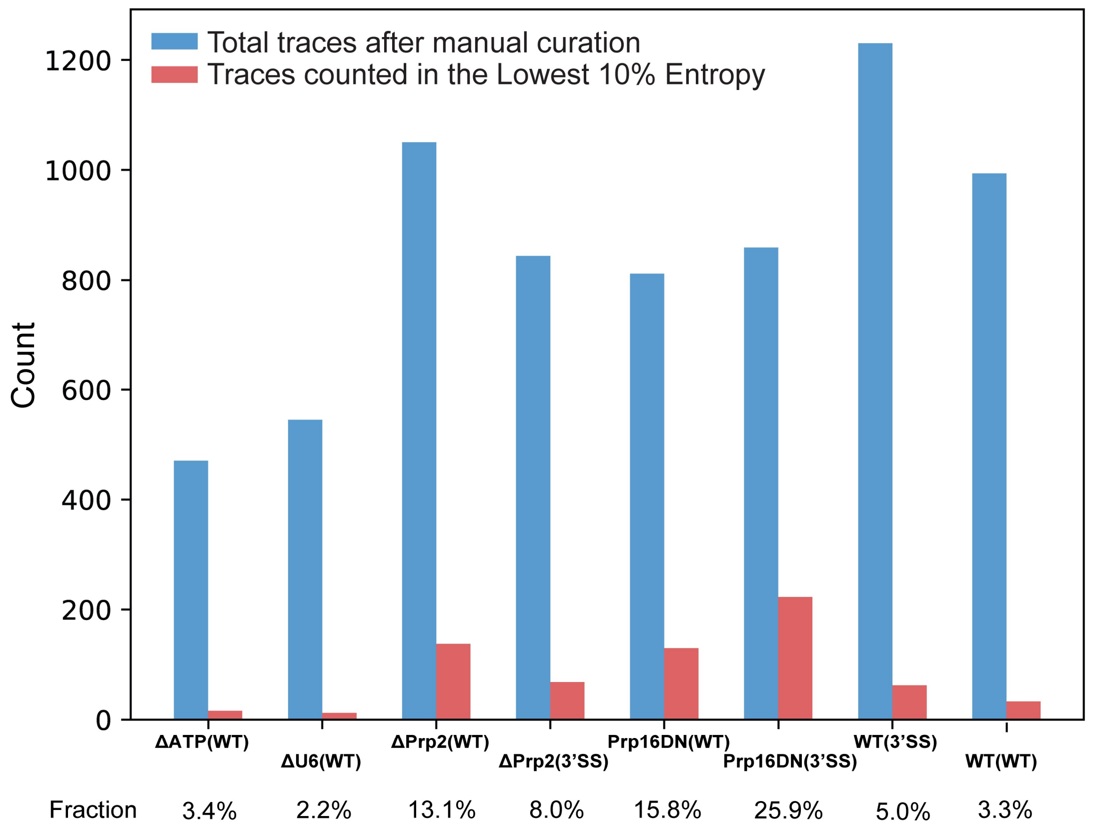


**Supplementary Fig. 8 | Distribution of the 10% of traces with lowest LSE across the different experimental conditions in the splicing study.** The total number count and fraction of traces from each experimental dataset that are among those with the 10% lowest LSE values is different, indicating that certain conditions exhibit a larger fraction of highly condition-specific traces.

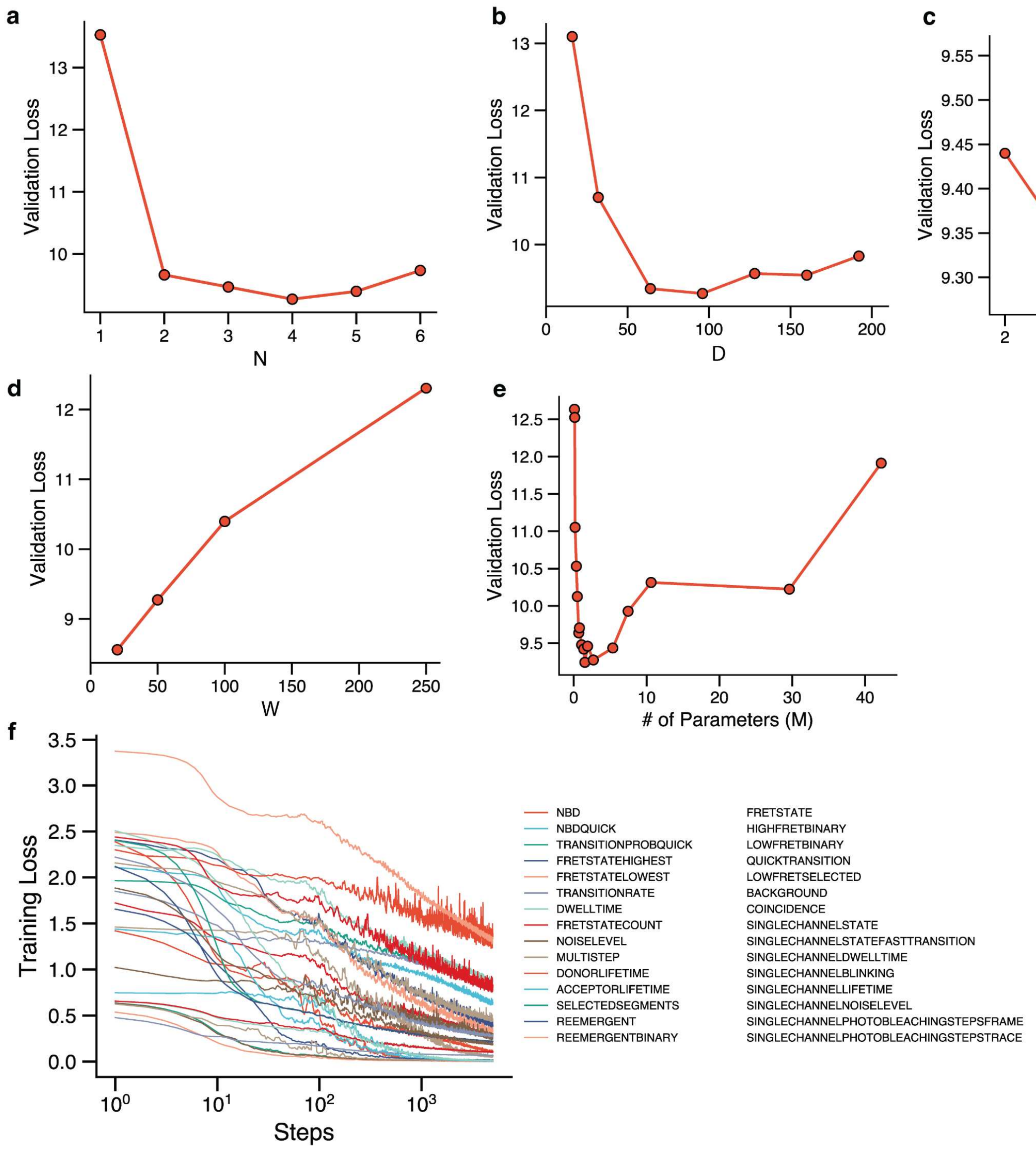


**Supplementary Fig. 9 | Pre-Training results and hyperparameter evaluation. a,** Impact of the number of transformer layers (N) on training effectiveness. **b,** Impact of the number of embedding vector dimensions (D) on training effectiveness. **c,** Impact of the number of Attention Heads on training effectiveness. **d,** Impact of the width of trace patches (W) used in tokenization on training effectiveness. **e,** Impact of the number of parameters (M) on training effectiveness. **f,** Loss of all training tasks as a function of pre-training steps.

**Supplementary Table 1.** Definition of training tasks for multi-task pre-training of META-SiM. The main data attribute for each task listed in the table was used for grouping the tasks for the principal projection and task alignment calculations in Fig. 4.

| Index | Task | Definition | Main Data Attribute | Frame-level or Trace-level |
|---|---|---|---|---|
| 1 | FRET state idealization | Predict true FRET state for each time frame. Prediction is FRET value ([0, 1]) instead of FRET state index (1, 2, 3, etc.). | Kinetic rate constant | Frame-level |
| 2 | Counting the number of FRET states | Counting the number of distinct FRET states presented in a time trace. | Number of states | Trace-level |
| 3 | Identifying highest FRET state frames | Locating which frames are in the FRET state with the highest FRET value. | FRET value | Frame-level |
| 4 | Identifying lowest FRET state frames | Locating which frames are in the FRET state with lowest FRET value, and label it as true. Note that blinking and photobleached states are not considered the lowest FRET state and labeled as false. | FRET value | Frame-level |
| 5 | Identifying highest FRET state's FRET value | Identifying the highest FRET value for a time trace. | FRET value | Trace-level |
| 6 | Identifying lowest FRET state's FRET value | Identifying the lowest FRET value that is not blinking or photobleached for a time trace. | FRET value | Trace-level |
| 7 | Extracting mean dwell time | Predicting the mean dwell time of all the FRET states in a time trace. | Kinetic rate constant | Trace-level |
| 8 | Extracting signal-to-noise ratio | Predicting the signal-to-noise ratio for a time trace. | Noise | Trace-level |
| 9 | Extracting donor fluorophore lifetime | Predicting the total lifetime of the donor fluorophore for a time trace. | FRET donor lifetime | Trace-level |
| 10 | Extracting acceptor fluorophore lifetime | Predicting the total lifetime of the acceptor fluorophore for a time trace. | FRET acceptor lifetime | Trace-level |
| 11 | Counting multistep photobleaching | Counting the photobleaching steps for traces with single- or multi-step photobleaching (0-4 steps). | Photobleaching steps | Trace-level |
| 12 | Selecting segments for active FRET states | Selecting the frames of a time trace where the FRET state is either static or dynamic, but is not blinking or photobleached. | Photobleaching steps | Frame-level |
| 13 | Extracting the average kinetic rate constant | Predicting the average kinetic rate constant for a | Kinetic rate constant | Trace-level |

| | | | | |
|---|---|---|---|---|
| | | time trace. The ground truth is defined as the harmonic average of all kinetic rate constants that are used in the simulator for a particular trace. | | |
| 14 | Identifying re-appearance of active FRET states after photobleaching | Identifying frames that are in active FRET states but occur after one or more photobleaching events. This behavior is common for experimental data where more than one fluorophore is present. | Photobleaching steps | Frame-level |
| 15 | Identifying existence of re-appearance of active FRET states after photobleaching | Identifying whether a trace has active FRET states after one or more photobleaching events. This behavior is common for experimental data. | Photobleaching steps | Trace-level |
| 16 | Counting the number of state transitions | Counting the total number of state transitions for a time trace. | Kinetic rate constant | Trace-level |
| 17 | Counting the number of state transitions for very fast dynamics | Counting the total number of state transitions for a time trace with very fast dynamics. | Kinetic rate constant | Trace-level |
| 18 | FRET state idealization for very fast dynamics | Predict true FRET state for each time frame with very fast dynamics. Prediction is FRET value ([0, 1]) instead of FRET state index (1, 2, 3, etc.) | Kinetic rate constant | Frame-level |
| 19 | Extracting average kinetic rate constant for very fast dynamics | Predicting the average kinetic rate constant for time traces with very fast dynamics. | Kinetic rate constant | Trace-level |
| 20 | Identifying if a frame is in the lowest FRET state | Identifying whether a frame in a time trace is in the FRET state with the lowest FRET value and is an active state. This task aims to help models distinguish real low FRET from photobleached states. Note that the previous prediction target of highest/lowest FRET state tasks is the FRET value, while this task's prediction target is a binary true/false label for each frame. | FRET value | Frame-level |
| 21 | Extracting background intensity | Predicting the true background intensity after the last photobleaching event for a time trace. | Noise | Trace-level |

| | | | | |
|---|---|---|---|---|
| 22 | Identifying colocalized pairs of fluorophores | Predicting whether a time trace has more than 1 pair of fluorophores. This phenomenon is common in experimental data when two or more pairs of fluorophores overlap in space (i.e., are colocalized). | Photobleaching steps | Trace-level |
| 23 | State idealization for a single-channel time trace | Identifying true states and predict de-noised intensity for a single-channel time trace. | Single-channel kinetic rate constant | Frame-level |
| 24 | Extract the mean dwell time for a single-channel time trace | Predicting the dwell time averaged over all kinetic states in a single-channel time trace. | Single-channel kinetics rate constant | Trace-level |
| 25 | Extract the average kinetic rate constant for a single-channel time trace | Predicting the average kinetic rate constant for a time trace. The ground truth is defined as the harmonic average of all kinetic rate constants that are used in the simulator for a particular single-channel trace. | Single-channel kinetics rate constant | Trace-level |
| 26 | Identifying blinking frames for a single-channel time trace | Identifying the frames that are in a blinking state for a single-channel time trace. | Single-channel noise | Frame-level |
| 27 | Extracting fluorophore lifetime for a single-channel time trace | Predicting the total lifetime of the fluorophore before the first photobleaching event for a single-channel time trace. | Single-channel lifetime | Trace-level |
| 28 | Extracting the signal-to-noise ratio for a single-channel time trace | Predicting the signal-to-noise ratio for a single-channel time trace. | Single-channel noise | Trace-level |
| 29 | Identifying photobleaching step frames for a single-channel time trace | Locating the frames where photobleaching steps happen for a single-channel time trace. | Single-channel photobleaching steps | Frame-level |
| 30 | Counting photobleaching steps for a single-channel time trace | Counting the total number of photobleaching steps for a single-channel time trace. | Single-channel photobleaching steps | Trace |

**Supplementary Table 2.** Description of datasets used in pre-training and downstream tasks.

| Dataset Index | Tasks | Description | Data Type | Pre-Training Set Size [Trace] | Fine-tuning Set Size [Trace] | Testing Set Size [Trace] | # of Detection Channels |
|---|---|---|---|---|---|---|---|
| D1 | Trace Classification & Segmentation | A toehold-exchange-based DNA walker[8] | Exp. | N/A | 109 | 642 | Two-color |
| D2 | | A DNA swinging arm[2] | | | 82 | 233 | |
| D3 | | A preQ$_1$ riboswitch[3] | | | 215 | 4628 | |
| D4 | | A paused transcriptional elongation complex[4] | | | 137 | 3770 | |
| D5 | | A $Mn^{2+}$ riboswitch[5] | | | 105 | 656 | |
| D6 | Stoichiometry Analysis | A photobleaching dataset from a 6-subunit protein complex bearing up to one HaloTag Alexa Fluor 660 (Alexa660) label per monomer | Exp. | N/A | 261 | 231 | One-color |
| D7 | Kinetic Fingerprinting | A use case involving detection of the *EGFR* point mutation T790M in DNA[6] | Exp. | N/A | 3904 (not manually curated*) | 5720 | One-color |
| D8 | Trace Idealization | ACTR-NCBD Binding at 10-ms binning[1,9] | Exp. | N/A | N/A** | 19*** | Two-color |
| D9 | Biological Discovery | Pre-mRNA conformational changes during yeast splicing[7] | Exp. | N/A | N/A | 6805 | Two-color |
| D10 | META-SiM Pre-Training | A synthetic dataset of smFRET traces representing diverse dynamics to train META-SiM | Simulated | 1.5 million | N/A | N/A | Two-color & One-color |
| D11 | smFRET Atlas Training & Annotation | A synthetic dataset of smFRET traces representing diverse dynamics to train and annotate a global 2D UMAP with META-SiM | Simulated | 1.02 million | N/A | N/A | Two-color |

* For the dataset D7, the labels for the fine-tuning set was not manually curated, but generated based on the experimental conditions, whereas true for mutant only condition and false for wild-type only condition, same strategy used in our previous work in ref[10].

** D8, D9 use cases didn't involve any fine-tuning. META-SiM foundation model was directly applied in these tasks.

*** The full experiment dataset used in the benchmark study of 14 tools[1], with n(traces) = 19, n(datapoints) = 226,100, using 10 ms time bins resulting in 100 Hz sampling.

**Supplementary Table 3.** Description and parameter range for generating traces to build the smFRET Atlas, 1 million traces for training and 22,000 traces for annotation. (All ranges in the table represent uniform distributions.)

| Code | Description | Number of States | SNR | FRET Values | Kinetic Rate Constant [Frame$^{-1}$] |
|---|---|---|---|---|---|
| 1-c-l | 1 FRET state, clean, low FRET value | 1 | [4, 8] | State 1: [0.05, 0.35] | N/A |
| 1-c-m | 1 FRET state, clean, middle FRET value | 1 | [4, 8] | State 1: [0.4, 0.6] | N/A |
| 1-c-h | 1 FRET state, clean, high FRET value | 1 | [4, 8] | State 1: [0.65, 0.95] | N/A |
| 1-n-l | 1 FRET state, noisy, low FRET value | 1 | [1.5, 4] | State 1: [0.05, 0.35] | N/A |
| 1-n-m | 1 FRET state, noisy, middle FRET value | 1 | [1.5, 4] | State 1: [0.4, 0.6] | N/A |
| 1-n-h | 1 FRET state, noisy, high FRET value | 1 | [1.5, 4] | State 1: [0.65, 0.95] | N/A |
| 2-c-lm-s | 2 FRET states, clean, middle and low FRET values, slow transition | 2 | [4, 8] | State 1: [0.05, 0.35]<br>State 2: [0.4, 0.6] | [0.005, 0.025] |
| 2-c-lm-f | 2 FRET states, clean, middle and low FRET values, fast transition | 2 | [4, 8] | State 1: [0.05, 0.35]<br>State 2: [0.4, 0.6] | [0.05, 0.2] |
| 2-c-lh-s | 2 FRET states, clean, low and high FRET values, slow transition | 2 | [4, 8] | State 1: [0.05, 0.35]<br>State 2: [0.65, 0.95] | [0.005, 0.025] |
| 2-c-lh-f | 2 FRET states, clean, low and high FRET values, fast transition | 2 | [4, 8] | State 1: [0.05, 0.35]<br>State 2: [0.65, 0.95] | [0.05, 0.2] |
| 2-c-mh-s | 2 FRET states, clean, middle and high FRET values, slow transition | 2 | [4, 8] | State 1: [0.4, 0.6]<br>State 2: [0.65, 0.95] | [0.005, 0.025] |
| 2-c-mh-f | 2 FRET states, clean, middle and high FRET values, fast transition | 2 | [4, 8] | State 1: [0.4, 0.6]<br>State 2: [0.65, 0.95] | [0.05, 0.2] |
| 2-n-lm-s | 2 FRET states, noisy, middle and low FRET values, slow transition | 2 | [1.5, 4] | State 1: [0.05, 0.35]<br>State 2: [0.4, 0.6] | [0.005, 0.025] |
| 2-n-lm-f | 2 FRET states, noisy, middle and low FRET values, fast transition | 2 | [1.5, 4] | State 1: [0.05, 0.35]<br>State 2: [0.4, 0.6] | [0.05, 0.2] |
| 2-n-lh-s | 2 FRET states, noisy, low and high FRET values, slow transition | 2 | [1.5, 4] | State 1: [0.05, 0.35]<br>State 2: [0.65, 0.95] | [0.005, 0.025] |
| 2-n-lh-f | 2 FRET states, noisy, low and high FRET values, fast transition | 2 | [1.5, 4] | State 1: [0.05, 0.35]<br>State 2: [0.65, 0.95] | [0.05, 0.2] |
| 2-n-mh-s | 2 FRET states, noisy, middle and high FRET values, slow transition | 2 | [1.5, 4] | State 1: [0.4, 0.6]<br>State 2: [0.65, 0.95] | [0.005, 0.025] |
| 2-n-mh-f | 2 FRET states, noisy, middle and high FRET values, fast transition | 2 | [1.5, 4] | State 1: [0.4, 0.6]<br>State 2: [0.65, 0.95] | [0.05, 0.2] |

| | | | | | |
|---|---|---|---|---|---|
| 3-c-lmh-s | 3 FRET states, clean, low and middle and high FRET values, slow transition | 3 | [4, 8] | State 1: [0.01, 0.35]<br>State 2: [0.4, 0.6]<br>State 3: [0.65, 0.95] | [0.002, 0.025] |
| 3-c-lmh-f | 3 FRET states, clean, low and middle and high FRET values, slow transition | 3 | [4, 8] | State 1: [0.01, 0.35]<br>State 2: [0.4, 0.6]<br>State 3: [0.65, 0.95] | [0.05, 0.2] |
| 3-n-lmh-s | 3 FRET states, noisy, low and middle and high FRET values, slow transition | 3 | [1.5, 4] | State 1: [0.01, 0.35]<br>State 2: [0.4, 0.6]<br>State 3: [0.65, 0.95] | [0.002, 0.025] |
| 3-n-lmh-f | 3 FRET states, noisy, low and middle and high FRET values, slow transition | 3 | [1.5, 4] | State 1: [0.01, 0.35]<br>State 2: [0.4, 0.6]<br>State 3: [0.65, 0.95] | [0.05, 0.2] |
